# Mitochondrial cyclophilin D promotes disease tolerance by licensing NK cell development and IL-22 production against influenza virus

**DOI:** 10.1101/2021.05.28.445832

**Authors:** Jeffrey Downey, Haley E. Randolph, Erwan Pernet, Kim A. Tran, Shabaana A. Khader, Irah L. King, Luis B. Barreiro, Maziar Divangahi

## Abstract

Immunity to infectious disease involves a combination of host resistance, which eliminates the pathogen, and disease tolerance, which limits tissue damage. While the severity of most pulmonary viral infections, including influenza A virus (IAV), is linked to excessive inflammation, our mechanistic understanding of this observation remains largely unknown. Here we show that mitochondrial cyclophilin D (CypD) protects against IAV infection via disease tolerance. Mice deficient in CypD (*CypD^-/-^* mice) are significantly more susceptible to IAV infection despite comparable antiviral immunity. Instead, this susceptibility resulted from damage to the lung epithelial barrier caused by a significant reduction of IL-22 production by conventional NK cells in IAV-infected *CypD^-/-^* mice. Transcriptomic and functional data revealed that the compromised IL-22 production by NK cells resulted from dysregulated lymphopoiesis, stemming from increased cell death in NK cell progenitors, as well as the generation of immature NK cells that exhibited altered mitochondrial metabolism. Importantly, following IAV infection, administration of recombinant IL-22 abrogated pulmonary damage and enhanced survival of *CypD^-/-^* mice. Collectively, these results demonstrate a key role for CypD in NK cell-mediated disease tolerance.

## INTRODUCTION

In response to any given infection, host resistance mechanisms are involved in preventing pathogen invasion or replication. These resistance mechanisms are a critical component of host defense to infection, yet they come with a considerable inflammatory cost that paradoxically threatens host fitness through excessive immunopathology. Thus, mechanisms of disease tolerance are required to mitigate tissue damage, restore organ function and counter the cost of anti-microbial inflammation (*1–4*).

Influenza A viruses (IAV) cause consistent and recurrent respiratory infections, responsible for approximately half a million deaths per annum globally, while also causing unpredictable and devastating pandemics. Severe and fatal IAV infections are more often triggered by a “cytokine storm” that contributes to a break in disease tolerance—characterized by destruction of the pulmonary epithelial/endothelial barrier and respiratory insufficiency—rather than ineffective host resistance (*5–7*). On the other hand, 16% of IAV infections are estimated to be asymptomatic (*8*) and similar observations have been made clinically during the COVID-19 pandemic. Like influenza, deaths due to COVID-19 seem to be more often associated with a loss of disease tolerance (*9, 10*), suggesting that in certain individuals these mechanisms are sufficient to avoid severe disease manifestation. Thus, understanding disease tolerance responses against a variety of acute respiratory infections may uncover targetable pathways for novel immunotherapies.

IAV infections are short-lived and usually self-resolving within approximately 7-10 days, though inflammation can persist for weeks after infection. The early inflammatory response is marked by an influx of innate leukocytes into the infected lung and airways. While inflammatory monocyte-derived cells play important roles in host resistance and priming of the adaptive immune response (*11–13*), they can jeopardize survival through immunopathology (*14, 15*). Mechanistically, our group recently highlighted the importance of the bioactive lipid leukotriene B_4_ (LTB_4_) in inhibiting *in situ* proliferation of inflammatory monocyte-derived macrophages (IMM) to conserve pulmonary epithelial integrity (*16*). Equally, reduction of extracellular matrix (ECM) turnover during IAV infection, by inhibiting lung metalloprotease activity, protected mice by conserving epithelial structure (*17*). Thus, maintenance of the pulmonary architecture and physiology is an essential component of host defense to IAV infection.

In addition to myeloid cells, innate lymphoid cells (ILCs), such as natural killer (NK) cells, accumulate in the lung shortly upon infection, beginning as early as 2 days post-infection (*18*). NK cells express a variety of activating and inhibitory receptors that perform diverse context- and tissue-specific functions, including well-described roles in the killing of virally-infected cells in a non-MHC-restricted manner through the production of IFN-γ and expression of perforin and granzymes (*19*). In the context of IAV infection, NK cells contribute to host resistance by recognizing sialylated hemagglutinin (HA) proteins (*20, 21*) on the surface of IAV-infected cells to facilitate lysis (*22–24*). Beyond their cytotoxic role in host resistance, NK cells are equally critical in disease tolerance by secreting the epithelium-protective cytokine IL-22 (*25*). IL-22 is a member of the IL-10 family of cytokines that maintains mucosal barriers by inducing survival and proliferation of epithelial cells. Although several cell types are capable of producing IL-22— including NKT cells, γδ T-cells, ILC3s and αβ T-cells (*26, 27*)—conventional NK cells have been suggested to be the major source following IAV infection (*28*), particularly at later time-points (*29*). Consistently, *Il22^-/-^* mice exhibit enhanced epithelial damage and pulmonary pathology in response to IAV (*26, 29, 30*). However, our understanding of the mechanisms involved in the production of IL-22 by NK cells in this setting is incomplete.

Mitochondrial Cyclophilin D (CypD), encoded in the nucleus by the *Peptidyl-prolyl isomerase F* (*Ppif*) gene, is a member of the cyclophilin family of isomerases that resides within the mitochondrial matrix. CypD is well-known as an essential modulator of the mitochondrial permeability transition pore (MPTP) which is required for the induction of necrosis (*31–33*). Given the regulatory role of CypD in macrophage necrosis (*32, 33*) and the importance of conserved macrophage viability/function in immunity to IAV (*34–36*), we initially hypothesized that the loss of CypD-mediated necrosis would enhance macrophage viability and, thus, protection to IAV infection. Surprisingly, CypD-deficient mice were highly susceptible to IAV infection without aberrations in host resistance mechanisms or leukocyte necrosis. A lack of IL-22 production by conventional NK cells in the infected airways of *CypD*^-/-^ mice was the major cause of susceptibility, as exogenous reconstitution of *CypD^-/-^* mice with IL-22, or transfer of wild type (WT), but not CypD-deficient, NK cells into *Il22^-/-^* mice significantly improved disease tolerance. RNA-seq of purified NK cells from IAV-infected mice revealed a unique transcriptome in CypD-deficient NK cells, marked by an immature profile (*37*). We found that CypD promotes NK cell responses at two levels: (1) by preventing cell death of NK cell progenitors in the bone marrow to facilitate NK cell hematopoiesis; and (2) by regulating peripheral NK cell metabolism, which is required for the conversion of immature to mature NK cells (*38*). Collectively, our findings highlight an essential role for CypD in NK cell-mediated disease tolerance to acute IAV infection.

## RESULTS

### CypD is required for host defense to influenza A virus infection by promoting disease tolerance

Given the requirement of CypD in necrosis (*31, 32*) and the relationship between influenza A virus (IAV) pathogenesis and cell death (*34*), we postulated that *CypD^-/-^* mice are more resistance to IAV infection. We first infected WT and *CypD^-/-^* mice with an LD_50_ (90 PFU) dose of PR8 IAV. *CypD^-/-^* mice were highly susceptible to IAV infection, exhibiting increased mortality (**Fig. 1A**) and morbidity (**Fig. 1B**). To delineate the cause of mortality, we hereafter used a sublethal (50 PFU) dose of IAV. Interestingly, the enhanced susceptibility of *CypD^-/-^* mice was not due to differences in pulmonary necrosis (**Fig. S1A**) or host resistance, as *CypD^-/-^* mice had similar pulmonary viral loads (**Fig. 1C**) and levels of active type I IFN (IFN-I) in the lung (**Fig. 1D-E; Fig. S1B-C**). We next hypothesized that the susceptibility of *CypD^-/-^* mice was linked to impaired disease tolerance responses, leading to increased lung tissue damage and pulmonary inflammation. Following IAV infection, CypD-deficient mice exhibited enhanced pulmonary edema (**Fig. 1F**) and barrier damage, as there was significant increase in protein and erythrocyte influx into the BAL (**Fig. 1G-H; Fig. S1D)**. Intranasal delivery of a fluorescently-labelled dextran molecule into infected mice further demonstrated a significant increase in lung epithelial/endothelial damage of *CypD*^-/-^ mice through a collective loss of fluorescence in the lung over time (**Fig. 1I**). Both flow cytometry and H&E staining showed that this increased pulmonary damage was associated with elevated inflammatory cells in the parenchyma and airways of *CypD^-/-^* lungs (**Fig. 1J-L**) beginning at 7 days post-infection, which coincided with the onset of enhanced tissue damage in these mice. Using Masson’s Trichrome stain, we equally observed enhanced collagen deposition and fibrosis in the lungs of *CypD^-/-^* mice (**Fig. 1M-N**). Taken together, these data show that CypD is required in immunity to IAV by regulating disease tolerance, rather than host resistance.

**Figure 1:**
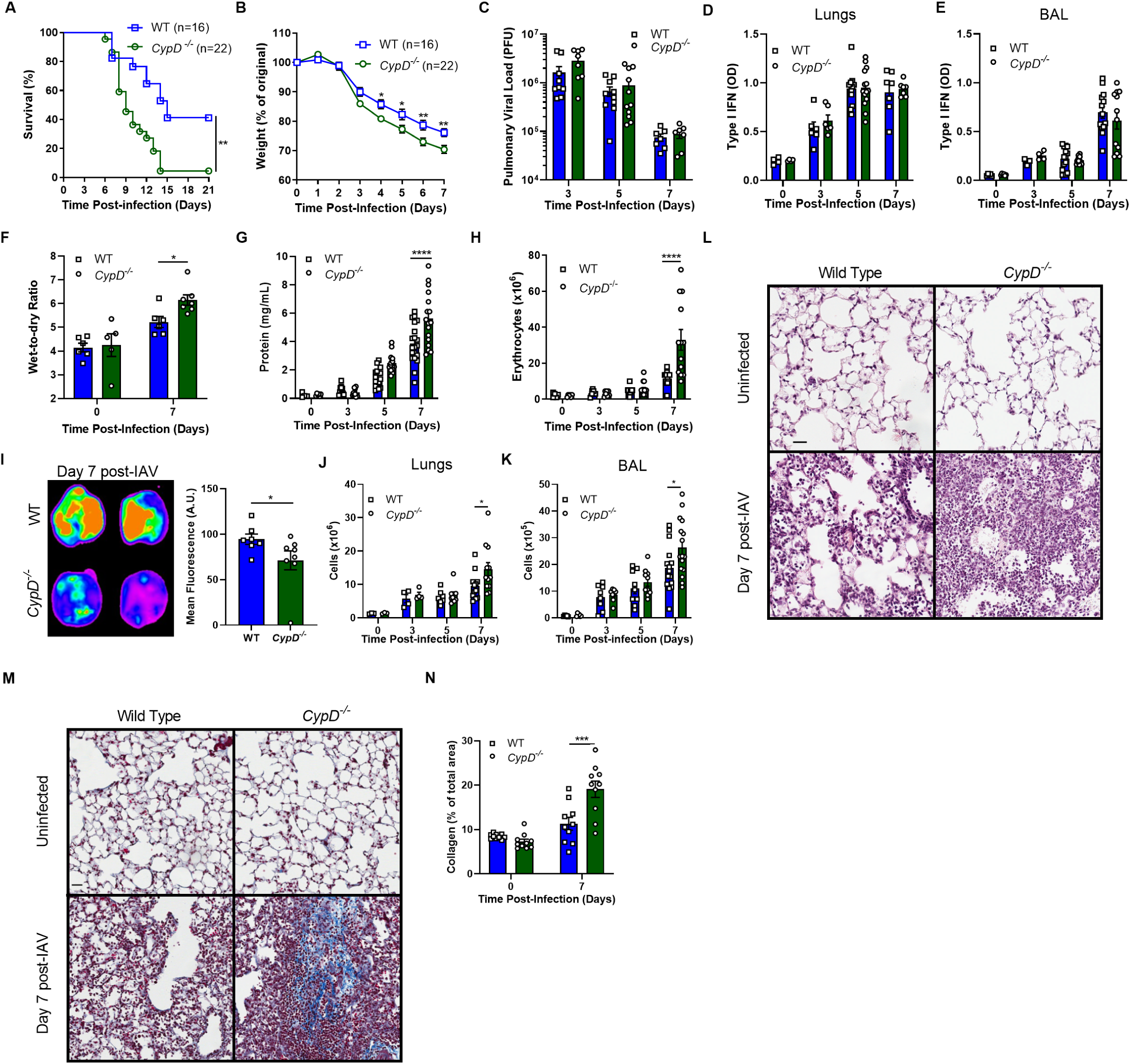
CypD protects against IAV infection by promoting disease tolerance. (A-B) WT and *CypD^-/-^* mice were infected with the LD_50_ dose of 90 PFU and survival (A) and weight loss (B) were monitored. (C-N) WT and *CypD^-/-^* mice were infected with a sublethal dose of 50 PFU. At various time points post-infection viral load (C), total active IFN-I in the lungs (D) or BAL (E) were measured. (F) Lung wet-to-dry ratio at steady-state or at day 7 post-infection. Protein (G) or erythrocytes (H) in the BAL of mice at various timepoints post-infection. (I) Fluorescence intensity of lungs following one hour of Texas-Red Dextran administration. The left panels are representative lung images from two separate experiments and are quantified on the right. Total cell counts in the lungs (J) and BAL (K) following infection. Representative micrographs of lungs stained with hematoxylin and eosin (L; scale bar = 30 µM) or Masson’s Trichrome (M; scale bar = 20µM) that are either uninfected or infected for 7 days, as quantified in (N). In A and B, total mice are denoted in the figures, in C-K each symbol represents a unique mouse, in L and M micrographs are representative of at least 5 mice and in N each symbol represents the quantification of one random micrograph (n=10 micrographs total). All figures are a compilation of at least two experiments. Statistical analyses were performed using the Log Rank Test (A), Two-way ANOVA followed by Sidak’s multiple comparison test (B-H, J-K, N) or Two-tailed Student’s T-Test (I). See also **Figure S1**.

### Immune, not stromal, cells are responsible for the susceptibility of *CypD^-/-^* mice to IAV

Mechanisms of disease tolerance can be mediated by either structural or hematopoietic compartments (*1, 3, 39*). To delineate which cellular compartment was primarily responsible for the increased tissue damage in *CypD^-/-^* mice, we generated bone marrow chimeric mice, where the hematopoietic compartment of lethally irradiated CD45.1 WT mice was reconstituted with CypD-deficient (CD45.2) (*CypD*^-/-^ → WT) bone marrow (BM) and vice versa (WT→ *CypD^-/-^*). After 10-12 weeks, reconstitution efficiency was greater than 92% (**Fig. S2A**). For further analysis, we chose day 7 post-IAV infection to match the peak lung damage and initial mortality of the CypD-deficient mice (**Fig. 1**). Chimeric mice reconstituted with *CypD*^-/-^ BM cells (*CypD*^-/-^ → WT) showed a statistically significant increase in erythrocytes in the BAL and enhanced pulmonary inflammation and collagen deposition when compared to WT (**Fig. 2A-D; Fig. S2B**). These results indicate that the reduced disease tolerance in CypD-deficient mice is predominately mediated by the hematopoietic compartment.

**Figure 2:**
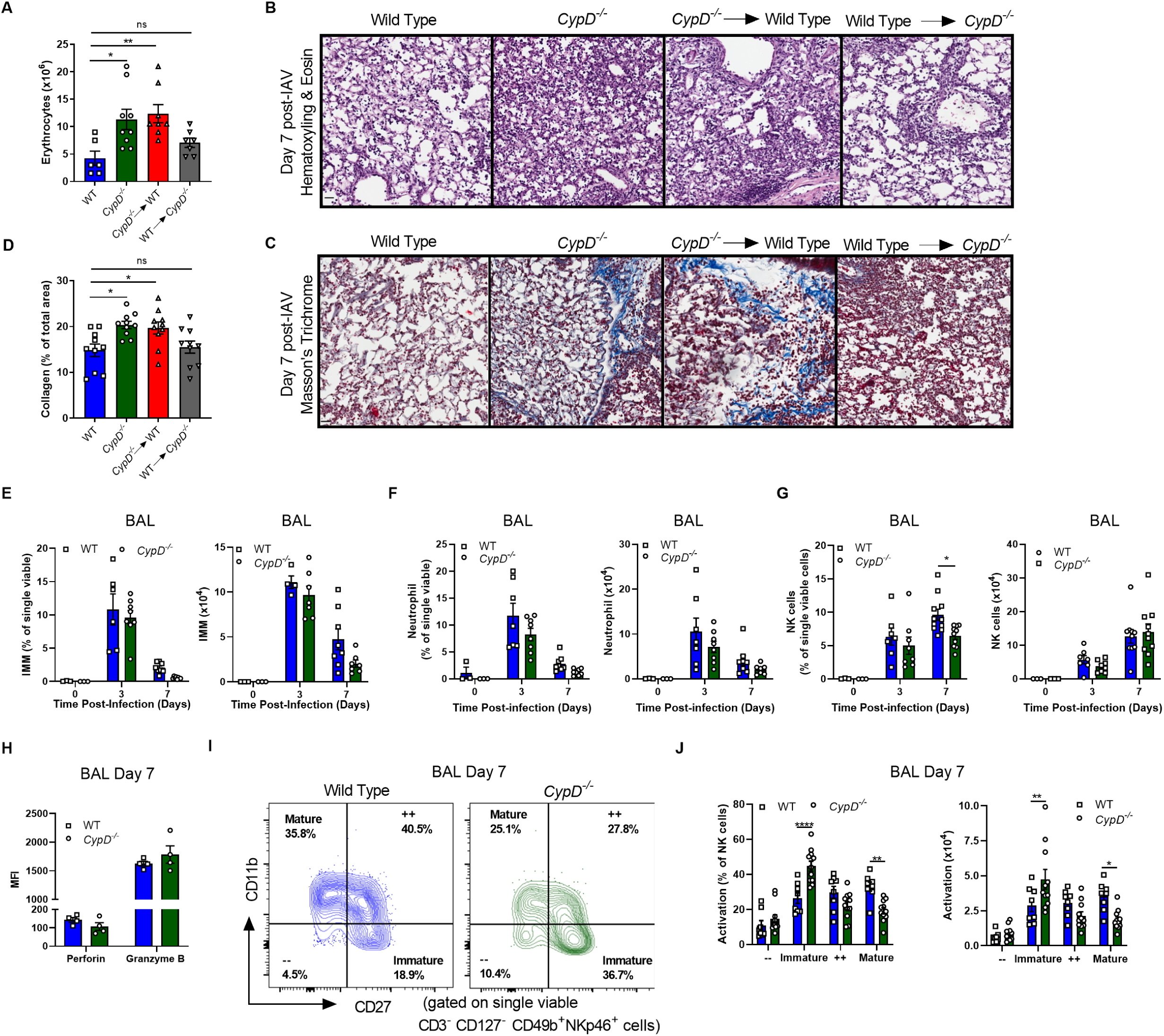
CypD mediates disease tolerance via hematopoietic cells and mice exhibit altered NK cell responses upon infection. (A-J) Mice were infected with 50 PFU of IAV. (A-D) Chimeric mice were generated by reconstituting irradiated CypD-deficient mice (CD45.2) with WT CD45.1 bone marrow (WT → *CypD^-/-^*) or irradiated CD45.1 mice with CypD-deficient (CD45.2) bone marrow (*CypD^-/-^* → WT). Chimeric mice were infected for 7 days and total erythrocytes in the BAL (A) were quantified. Representative micrographs of hematoxylin and eosin-stained lungs (B; scale bar = 30 µM) or Masson’s Trichrome (C; scale bar = 30 µM) as quantified in (D). (E-G) BAL from WT and *CypD^-/-^* mice were phenotyped by flow cytometry and total frequencies (left panels) and total cell counts (right panels) of IMM (E), neutrophils (F) and NK cells (G) enumerated. Mean fluorescence intensities of perforin and granzyme B of NK cells at day 7 (H) post-IAV infection. (I) Representative FACS plot of CD27 and CD11b expression on NK cells in the BAL of WT versus CypD-deficient mice at 7 days post-infection. (J) Quantifications of the percentages (left panel) and total cells counts (right panel) of NK cell activation subsets in the BAL at day 7 post-infection. In each panel, each symbol indicates a separate mouse, except in B-C, which is a representative figure of 4 mice/group, D, where each symbol is a randomly quantified micrograph (n=10 micrographs) and I, which is a representative plot compiled in J. All figures are a compilation of at least two experiments, except for H, which is one representative experiment of two. Statistical analyses were performed using the One-way ANOVA followed by Tukey’s multiple comparisons (A, D), Two-way ANOVA followed by Sidak’s multiple comparison test (E-G, J) or Two-tailed Student’s T-Test (H). See also **Figure S2**.

To elucidate which cell(s) of the hematopoietic compartment is/are responsible for the increased lung damage, we extensively phenotyped innate immune cells by flow cytometry (**full gating strategy Fig. S1 E-F**) at various time points post-IAV infection in the lung parenchyma and airways. We and others have previously shown that elevated levels of IMM and neutrophils can compromise disease tolerance during IAV infection (*11, 16*). However, the kinetics of IMM and neutrophils in the BAL showed no significant differences in frequencies or numbers between WT and *CypD^-/-^* mice (**Fig. 2E-F**) and even showed a trending decrease in *CypD^-/-^* mice. Interestingly, we found that the kinetics of NK cell accumulation in the airways differed between CypD-deficient and WT mice, with a significant reduction in the frequency of NK cells at day 7 post-IAV infection in *CypD^-/-^* mice, which aligned with the pulmonary damage (**Fig. 2G**). No differences in any of the cell populations assessed were observed in the lung parenchyma (**Fig. S2C-E**), hinting at the importance of spatial lung immunity (i.e. parenchyma versus airways) during IAV infection.

### CypD-deficient NK cells are phenotypically immature

NK cells are required for protection against IAV by regulating both host resistance and disease tolerance (*23, 29*). Correspondingly, a recent study suggested that altered kinetics of NK cells may underlie individual susceptibility to IAV, without affecting early viral loads (*40*). Thus, we next sought to further interrogate the role of NK cells in the susceptibility of CypD-deficient mice to IAV. NKp46/IAV HA interactions elicit cell lysis via perforin and granzyme to kill virally infected cells (*21, 23*). However, CypD-deficient and WT NK cells expressed similar levels of perforin and granzyme B (**Fig. 2H**), suggesting the lytic capacity of CypD-deficient NK cells was intact, as predicted given the similar viral loads in these mice. The maturation and accumulation of effector functions in NK cells is known to be dependent on downregulation of CD27 and upregulation of CD11b, which gives rise to four distinct populations of NK cells (CD27^-^ CD11b^-^, CD27^+^ CD11b^-^, CD27^+^ CD11b^+^, CD27^-^ CD11b^+^) (*37, 41, 42*). Unbiased single cell analysis has confirmed the existence of these populations and their importance in determining effector function of NK cells in mice (*43*). Intriguingly, *CypD^-/-^* NK cells exhibited a significantly higher frequency and number of immature (CD27^+^ CD11b^-^) cells compared to fully mature (CD27^-^ CD11b^+^) cells in the BAL (**Fig. 2I-J**), but not the lung (**Fig. S2F-G**) at day 7 post-IAV infection. Similar results as in the BAL were obtained in splenic NK cells at day 5 post-IAV infection (**Fig. S2H**). Collectively, our results show that CypD-deficient mice have reduced accumulation of mature NK cells in the airways at the peak of IAV-induced immunopathology.

### *CypD^-/-^* NK cells have a unique transcriptional profile and metabolic program following IAV infection

As NK cell kinetics and maturity differed between WT and *CypD^-/-^* mice following IAV infection, we next asked whether *CypD* was expressed in NK cells. To investigate this, we purified NK cells from WT and CypD-deficient spleens and found significant expression of *CypD* in WT, but not *CypD^-/-^* cells, both at steady-state and upon infection (**Fig. S3A**), which aligned with data publicly available from ImmGen (*44*). We then performed bulk RNA-seq on WT and CypD-deficient splenic NK cells isolated at day 5 post-IAV infection (**Table S1**). We chose day 5 post-infection, as splenic *CypD^-/-^* NK cells already exhibited an immature effector state when compared to WT (**Fig. S2H**). Principal component analysis (PCA) on the splenic WT and *CypD^-/-^* NK cell expression data revealed a strong signature of IAV infection, with non-infected and IAV-infected samples separating on PC1, which explains 56.3% of the variance in the dataset (**Fig. S3B**). Within the IAV-infected samples, we found 315 genes for which expression patterns significantly differed (|logFC| > 0.5, FDR < 0.10) between WT and *CypD^-/-^* mice (**Fig. 3A; Table S2**), including *Ppif* in line with our qPCR data (**Fig. S3A**). Gene ontology (GO) enrichment analysis revealed that genes showing significantly higher expression in the WT mice (n = 146) were enriched for pathways related to activation of the immune response (FDR = 4.3×10^-4^), phagocytosis (FDR = 5.3×10^-5^), wound healing (FDR = 4.1×10^-4^), and blood coagulation (FDR = 5.3×10^-4^) (**Fig. 3B; Table S3**), while genes more highly expressed in the *CypD^-/-^* mice (n = 169) were enriched for oxidative phosphorylation (OXPHOS) (FDR = 2.5×10^-3^), T cell mediated toxicity (FDR = 3.2×10^-3^), porphyrin-containing compound metabolic processes (FDR = 5.1×10^-3^), and peroxidase activity (FDR = 1.5×10^-4^) (**Fig. 3C; Table S3**). In line with our observation that *CypD^-/-^* mice display significantly more immature (CD27^+^CD11b^-^) and fewer fully mature (CD27^-^ CD11b^+^) NK cells, we found that genes previously associated with elevated expression in fully mature and intermediate-mature (CD27^+^ CD11b^+^) NK cells (Chiossone et al., 2009) were significantly enriched among genes showing higher expression in WT compared to *CypD^-/-^* mice (**Fig. S3C**) and that key genes driving these enrichments exhibited lower expression in *CypD^-/-^* mice (**Fig. 3D**).

**Figure 3:**
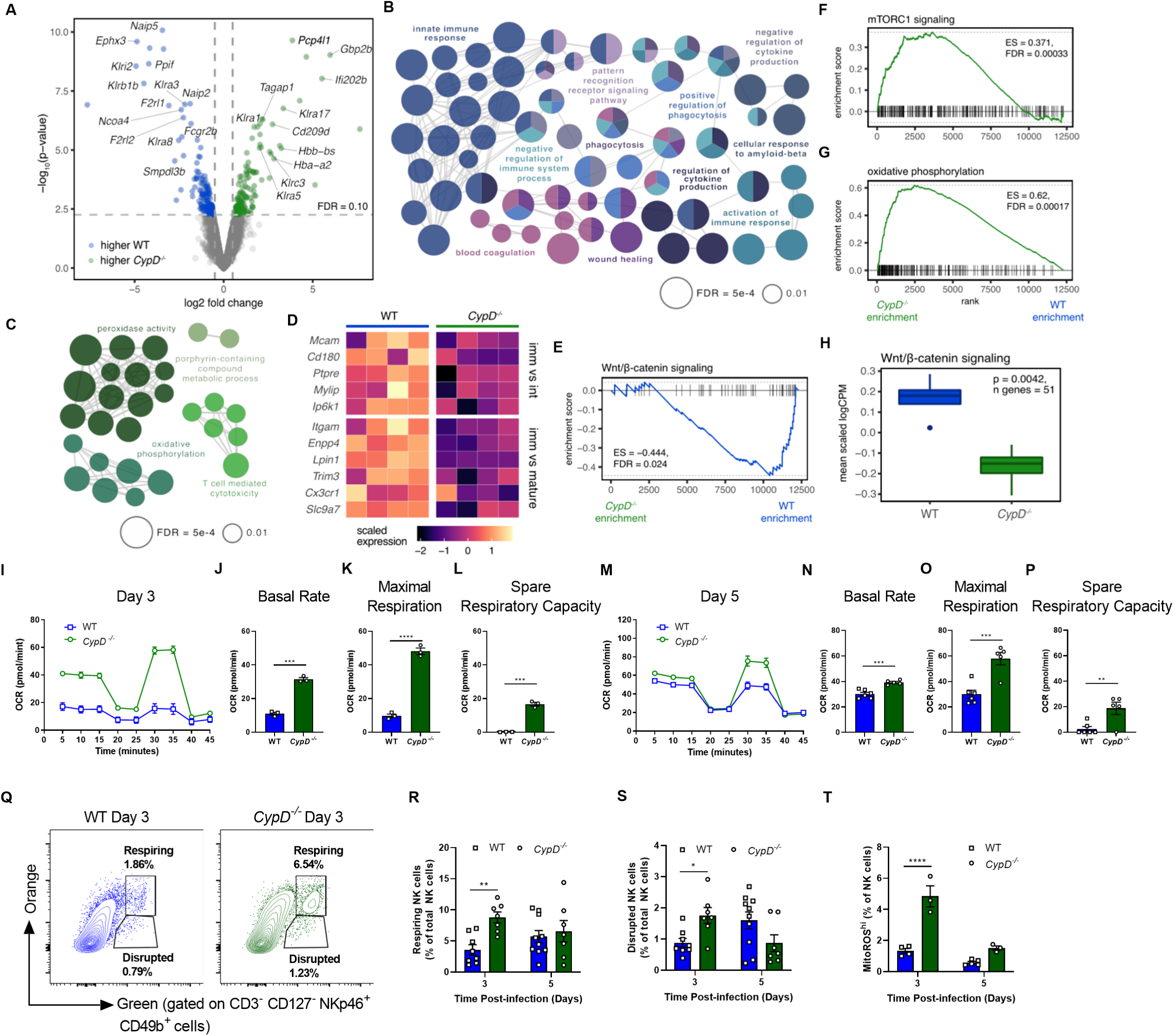
*CypD^-/-^* alters the transcriptome of IAV-infected splenic NK cells and induces a distinct metabolic phenotype. (A) Volcano plot of genes significantly differentially expressed (DE) between IAV-infected splenic NK cells of WT (blue, log2 fold change (FC) < -0.5, FDR < 0.10) and *CypD^-/-^* (green, log2 FC > 0.5, FDR < 0.10) mice. Genes with a negative log2 FC show higher expression in WT mice, while genes with a positive log2 FC show higher expression in *CypD^-/-^* mice. Selected genes show the most extreme changes in expression between the two genotype groups. (B) Significant (FDR < 0.01) ClueGO pathway enrichments for genes showing higher expression (FDR < 0.10) in the WT mice versus (C) *CypD^-/-^* mice in IAV-infected splenic NK cells. (D) Heatmap showing expression levels (mean-centered and scaled) for a subset of maturity marker genes as identified in Chiossone et al. (*37*) in IAV-infected NK cells of WT and *CypD^-/-^* mice. (“imm” = immature NK [CD27+CD11b-], “int” = intermediate-mature NK [CD27+CD11b+], “mature” = mature NK [CD27-CD11b+]). (E) Barcode enrichment plots for the hallmark Wnt/β-catenin signaling, (F) mTORC1 signaling and (G) oxidative phosphorylation pathways. A positive enrichment score (ES) corresponds to pathway enrichment among genes more highly expressed in *CypD^-/-^* mice (green), while a negative ES corresponds to pathway enrichment among genes more highly expressed in WT mice (blue). (H) Average, scaled logCPM expression estimates across genes in the hallmark Wnt/β-catenin signaling pathway. Oxygen consumption on purified splenic NK cells at day 3 (I-L) or day 5 (M-P) post-infection. (I, M) Representative Seahorse curves from which the values in (J-L) and (N-P) were calculated, respectively. (Q) Representative flow cytometry plots of splenic NK cells differentially stained for Mitotracker Orange and Mitotracker Green, as quantified in (R-S). Healthy respiring mitochondria were considered as Orange^hi^ and Green^hi^ (R) and disrupted mitochondria were considered as Orange^lo^ and Green^hi^ (S). (T) Frequency of MitoSox Red^hi^ NK cells in the spleen. (I-P) All data are from one representative experiment of two independent experiments and each dot is indicative of a technical replicate of NK cells compiled from 5 different mice. (Q-T) Data are compiled from two independent experiments and each dot represents a unique mouse. Statistical differences were assessed by Student’s T-Test (J-L and N-P) or Two-way ANOVA followed by Sidak’s multiple comparisons test (R-T). See also **Figure S3**.

Gene set enrichment analysis (GSEA) using the Molecular Signatures Database hallmark gene sets (*45*) highlighted a divergence in multiple metabolic and signaling pathways between WT and *CypD^-/-^* mice, including an enrichment for Wnt/β-catenin signaling among genes more highly-expressed in WT mice (FDR = 0.024) (**Fig. 3E; Table S3**), and enrichments for mTORC1 signaling (FDR = 3.3×10^-4^) and OXPHOS (FDR = 1.7×10^-4^) among genes more highly-expressed in *CypD^-/-^* mice (**Fig. 3F-G; Table S3**), suggesting a distinct metabolic program in CypD-deficient NK cells. Concordant with our enrichment analyses, the average expression of genes belonging to the Wnt/β-catenin signaling pathway was significantly higher in WT compared to *CypD^-/-^* mice (**Fig. 3H**, t-test, p = 4.2×10^-3^), while for the OXPHOS and mTORC1 signaling pathways, we observed a trend toward higher expression in the *CypD^-/-^* compared to WT mice (**Fig. S3D-E**, t-test, p = 0.06 and p = 0.24, respectively).

The RNA-seq results underscored an immature effector transcriptome in CypD-deficient NK cells, including decreased expression of the Wnt/β-catenin signaling pathway that has been shown to be a critical in the progression to mature CD11b^+^ NK cells (*46*), as well as an upregulation of OXPHOS and mTORC1 signaling, both of which have been implicated in immature effector states and impaired activation of NK cells (*38, 47, 48*). CypD is equally known to regulate ATP synthase and OXPHOS through incompletely understood mechanisms (*49*). To see whether an enhanced OXPHOS pathway expression led to elevated mitochondrial respiration in CypD-deficient NK cells, we purified splenic NK cells and subjected them to the Seahorse assay. At steady-state, we detected no differences in OXPHOS in WT and *CypD^-/-^* NK cells (**Fig. S3F-I**). However, over the course of infection, we observed a significant increase in basal metabolic rate, maximal respiration and spare respiratory capacity in *CypD^-/-^* NK cells compared to WT (**Fig. 3I-P**). We also found an increase in healthy respiring and disrupted mitochondria in CypD-deficient NK cells at day 3 post-infection (**Fig. 3Q-S**), as well as enhanced mitochondrial ROS production **(Fig. 3T**), suggesting the elevated OXPHOS contributes to mitochondrial dysregulation. Taken together, RNA-seq, flow cytometry and functional data indicate that CypD within NK cells enables their maturation and regulates cellular metabolism.

### *CypD* expression in NK cells promotes transcriptional pathways involved in wound healing and hemostasis

We next sought to dissect the transcriptional differences in functional immune-related pathways between WT and CypD-deficient NK cells following IAV infection. Numerous killer cell lectin-like receptor genes (*Klr*) were significantly differentially-expressed between NK cells of WT and *CypD^-/-^* mice post-infection (**Fig. 3A**), including *Klri2*, *Klr8*, *Klra17*, and *Klra5*, among others, suggesting that the inhibitory and activating capacity of these NK cells may differ, which in turn may alter their functional capabilities. Indeed, the transcriptional immune response signatures of IAV-infected WT and *CypD^-/-^* NK cells were distinct from one another: genes displaying significantly higher expression in *CypD^-/-^* NK cells were enriched for antigen processing (FDR = 7.8×10^-3^) and T cell mediated cytotoxicity (FDR = 3.2×10^-3^) pathways (**Fig. 4A; Table S3**), whereas genes more highly-expressed in WT NK cells were enriched for effector functions related to phagocytosis and engulfment (FDR = 1.1×10^-3^) and complement-dependent cytotoxicity (FDR = 8.0×10^-4^) (**Fig. 4B; Table S3**). Notably, among genes more highly-expressed in WT NK cells, we also identified significant enrichments for multiple pathways involved in cytokine secretion, including IL-2 production (FDR = 8.1×10^-3^), IL-10 secretion (FDR = 1.2×10^-3^) and production (FDR = 5.6×10^-3^), negative regulation of IL-13 secretion (FDR = 6.8×10^-4^), and chemokine secretion (FDR = 1.8×10^-3^), (**Fig. 4B; Table S3**), suggesting that *CypD^-/-^* NK cells display impaired anti-inflammatory cytokine production compared to WT NK cells. Finally, in agreement with our previous observation that the WT NK cell transcriptome was enriched for pathways involved in blood coagulation (**Fig. 3B**) and that CypD-deficient mice had enhanced hemorrhaging (**Fig. 1H; Fig. S1D**), genes upregulated in WT compared to *CypD^-/-^* cells were enriched for hemostasis (FDR = 5.5×10^-4^). Moreover, negative regulation of fibroblast growth factor production (FDR = 1.5×10^-3^) was enriched in WT NK cells, which might explain the reduced fibrosis in WT lungs following IAV infection (**Fig. 1M-N**). Interestingly, one gene significantly upregulated in this pathway in WT cells, *Cd59a*, has previously been shown to mediate protection against IAV, in part, via inhibiting pulmonary hemorrhaging and fibrosis in the lung (*50*), suggesting a complementary role of CypD. Finally, genes showing significantly higher expression in WT mice were more likely to be found in the tissue repair gene set (Yanai et al., 2016) than expected by chance (WT p = 0.009), while no such enrichment was observed among genes more highly-expressed in *CypD^-/-^* mice (*CypD^-/-^* p = 0.798) (**Fig. 4C**), further underscoring the role of CypD in mediating disease tolerance.

**Figure 4:**
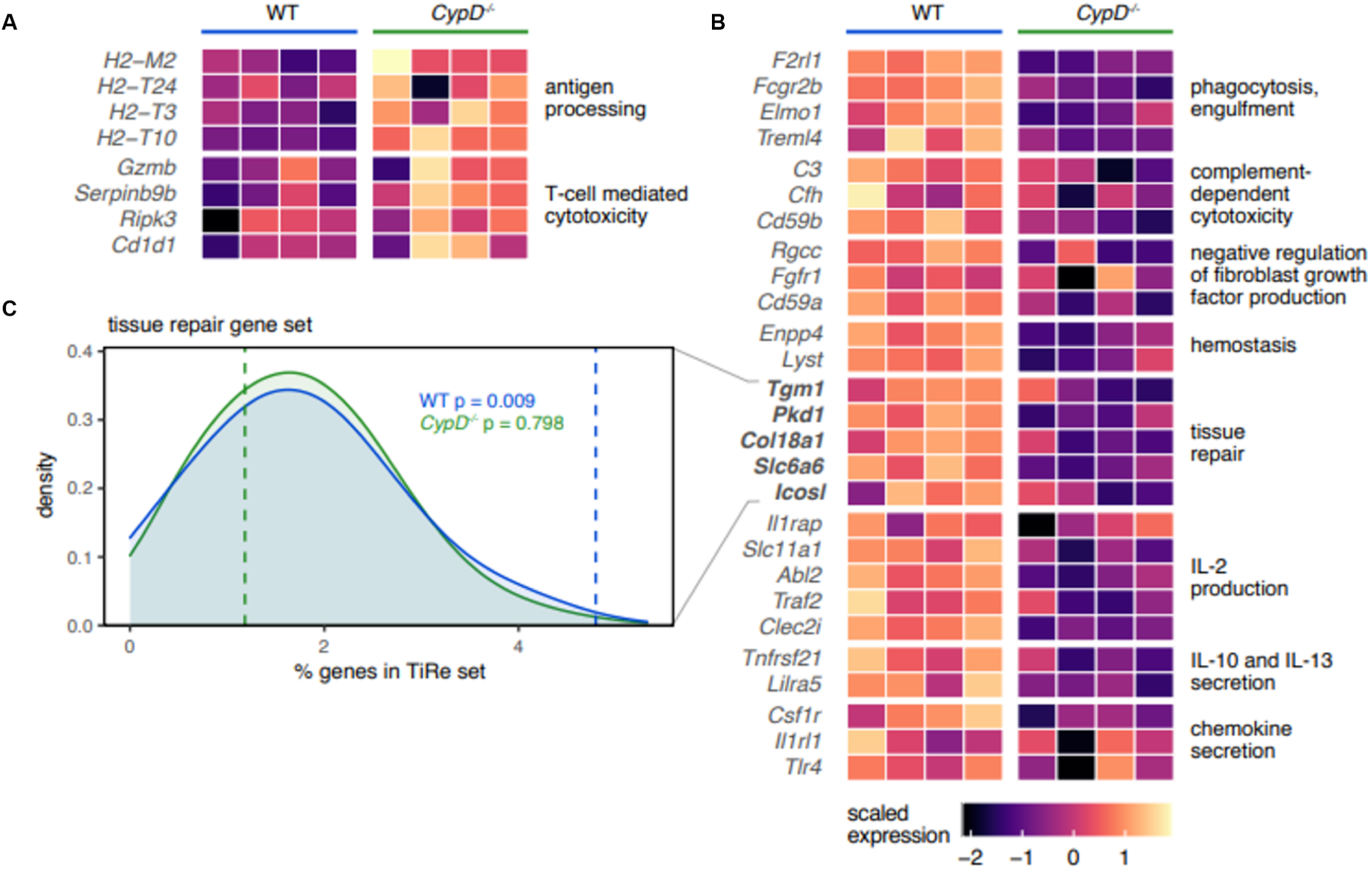
Splenic NK cells of *CypD^-/-^* mice are functionally reprogrammed. (A) Heatmap showing expression levels (mean-centered and scaled) for selected subsets of genes that fall into ontology pathways and functional gene sets significantly enriched among WT or (B) *CypD^-/-^* gene sets in IAV-infected splenic NK cells. (C) Proportion of WT or *CypD^-/-^* differentially-expressed genes that are in the tissue repair (TiRe) dataset (*98*) (WT, blue dashed line, p = 0.009; *CypD^-/-^*, green dashed line, p = 0.798) compared to random expectation when sampling the same total number of WT or *CypD^-/-^* differentially-expressed genes 1,000 times from all genes tested (null, density distributions).

### CypD promotes NK cell hematopoiesis within bone marrow progenitors to generate mature peripheral NK cells

Considering the functional role of CypD in cell death, the lack of developmentally mature NK cells in the airways of *CypD^-/-^* mice following IAV infection might be due to perturbations in local proliferation and/or cell death. To investigate these possibilities, we analyzed markers of proliferation (Ki67) and cell death (active caspase 3) by flow cytometry within NK cells in the BAL and found no differences (**Fig. 5A-B**). In addition to proliferation and cell death, aberrant recruitment or chemotaxis into the airways could explain the reduced number of mature NK cells in CypD-deficient mice. CCR2 expression on NK cells is known to specifically facilitate migration of NK cells into the airways, without affecting extravasation into the lung, which is instead mediated by CX3CR3 (*18, 51*). However, the frequency of CCR2^+^ NK cells was indistinguishable between WT and *CypD^-/-^* mice in the BAL, lung and blood (**Fig. 5C; Fig. S4A-B**), confirming CCR2-mediated migration of NK cells into the airways is not dependent on CypD expression.

**Figure 5:**
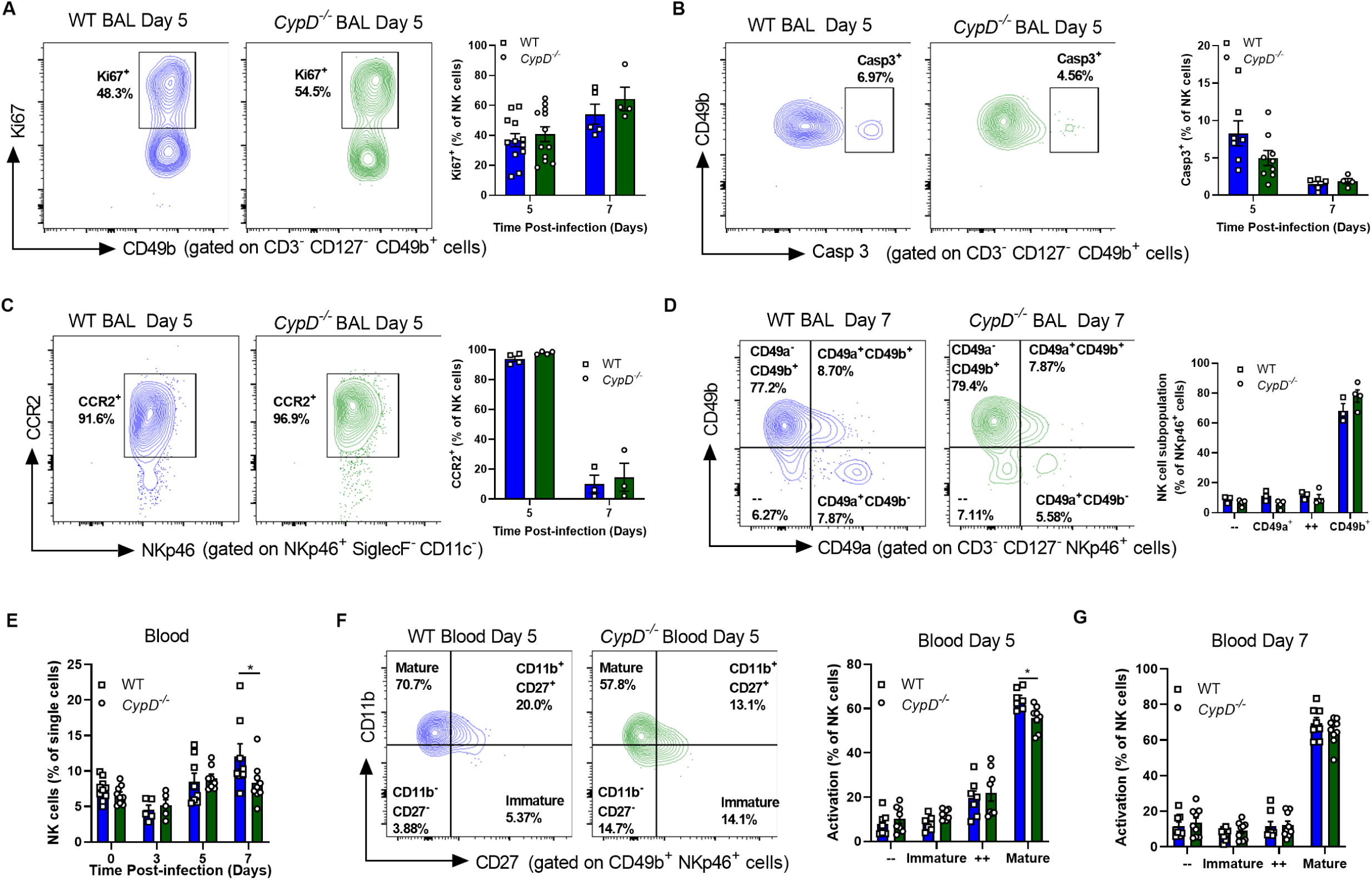
CypD-deficient mice have impaired recruitment of activated NK cells from the peripheral blood. (A-G) WT and *CypD^-/-^* were infected with 50 PFU. (A) Intracellular expression of Ki67, indicative of proliferative cells, in NK cells of the BAL. Left panels show a representative FACS plot as quantified on the right at day 5 post-infection. (B) Intracellular expression of active Caspase 3 expression by flow cytometry, indicative of apoptotic NK cells in the BAL of infected mice. The panels on the left are of a representative FACS plot at day 5 post-infection, as quantified on the right. (C) CCR2 expression on NK cells of the BAL with a day 5 representative FACS plot on the left and quantified on the right. (D) Differential expression of CD49b versus CD49a on NK cells of the BAL with a representative FACS plot on the left. (E-F) At various times post-infection peripheral blood was collected and the frequency of NK cells (E) and their activation state as defined by CD27 and CD11b expression (F-G) were quantified. In the left panel of F is a representative FACS plot of NK cells at day 5 post-infection. Each panel represents the compilation of at least two experiments, except C which is one representative experiment of two independent experiments. Each symbol represents one unique mouse. For all panels, Two-way ANOVA followed by Sidak’s multiple comparisons test was performed to determine significance. See also **Figure S4**.

Although continual on-demand egress of NK cells from the BM into the blood to supply peripheral tissues is well-described, it is now known that an additional population of long-lived tissue-resident NK cells exists in several peripheral tissues that can be differentiated by expression of CD49a rather than CD49b (*52*). This population is most prevalent in the uterus, liver and skin, but can be found in a variety of other sites, including the lung (*53*). To see whether this population of tissue-resident NK cells was altered in our model, we investigated expression of CD49a versus CD49b on the surface of NK cells from WT and *CypD^-/-^* mice. The vast majority of NK cells from the BAL and lungs of both groups of mice were CD49b^+^ CD49a^-^, indicative of a predominantly blood-derived population, and this frequency was the same between both groups (**Fig. 5D; Fig. S4C**). Thus, differences in blood-derived versus resident NK cells is not the reason for the altered accumulation and effector states observed in *CypD^-/-^* mice. Interestingly, the frequency of NK cells in the peripheral blood was decreased in CypD-deficient mice (**Fig. 5E**), and there were less fully mature CD27^-^ CD11b^+^ cells at day 5 post-infection, but not at other time points (**Fig. 5F-G; Fig. S4D-E**), suggesting NK cell recruitment was altered in *CypD^-/-^*mice.

The reduction of fully mature NK cells in the blood and peripheral tissue led us to speculate that there was a defect in the generation of NK cells in the BM of *CypD^-/-^* mice. NK cell generation occurs through a stepwise progression of progenitors, ultimately giving rise to fully mature effector NK cells that are released into the bloodstream (**Fig. 6A**). Downstream of the pluripotent LKS population, cells of the lymphoid lineage and the myeloid/erythroid lineage separate through the common lymphoid progenitor (CLP; Lin^-^ CD127^+^ cKit^lo^ Sca1^lo^) and the common myeloid progenitor (CMP; Lin^-^ CD127^-^ cKit^+^ Sca1^-^ CD34^+^ CD16/32^-^), respectively. Deriving from the CLP, NK cells are generated first from the pre-NK progenitor (pre-NKP; Lin^-^ CD127^+^ CD27^+^ CD122^-^ CD244.2^+^), then the NKP (Lin^-^ CD127^+^ CD27^+^ CD122^+^ CD244.2^+^), which represents what is thought to be the first fully committed cell of the NK lineage, and finally *bona fide* NK cells that then begin to express NKp46, CD49b and CD11b (*54*). To investigate any potential aberrations in NK cell development between WT and *CypD^-/-^* mice, we phenotyped the BM precursors beginning at the LKS population and working towards more committed progenitors, during homeostasis and upon infection. We observed no differences in the frequency or number of LKS cells between groups (**Fig. 6B; Fig. S4F**), nor in the CLP (**Fig. 6C; Fig. S4G**). Additionally, there were similar levels of the CMP and granulocyte-monocyte progenitor (GMP) in WT and CypD-deficient mice (**Fig. S4H-I**). However, beginning at the NK cell-specific progenitors, we observed a decrease in pre-NKP and NKP populations (**Fig. 6D-F**), as well as a lower frequency of fully mature CD11b^+^ CD27^-^ expressing NK cells in the BM upon infection (**Fig. 6G**; **Fig. S4J**). Thus, CypD mediates NK cell hematopoiesis and an inability to progress through the NK cell lineage correlates with a lack of mature NK cells in the periphery.

**Figure 6:**
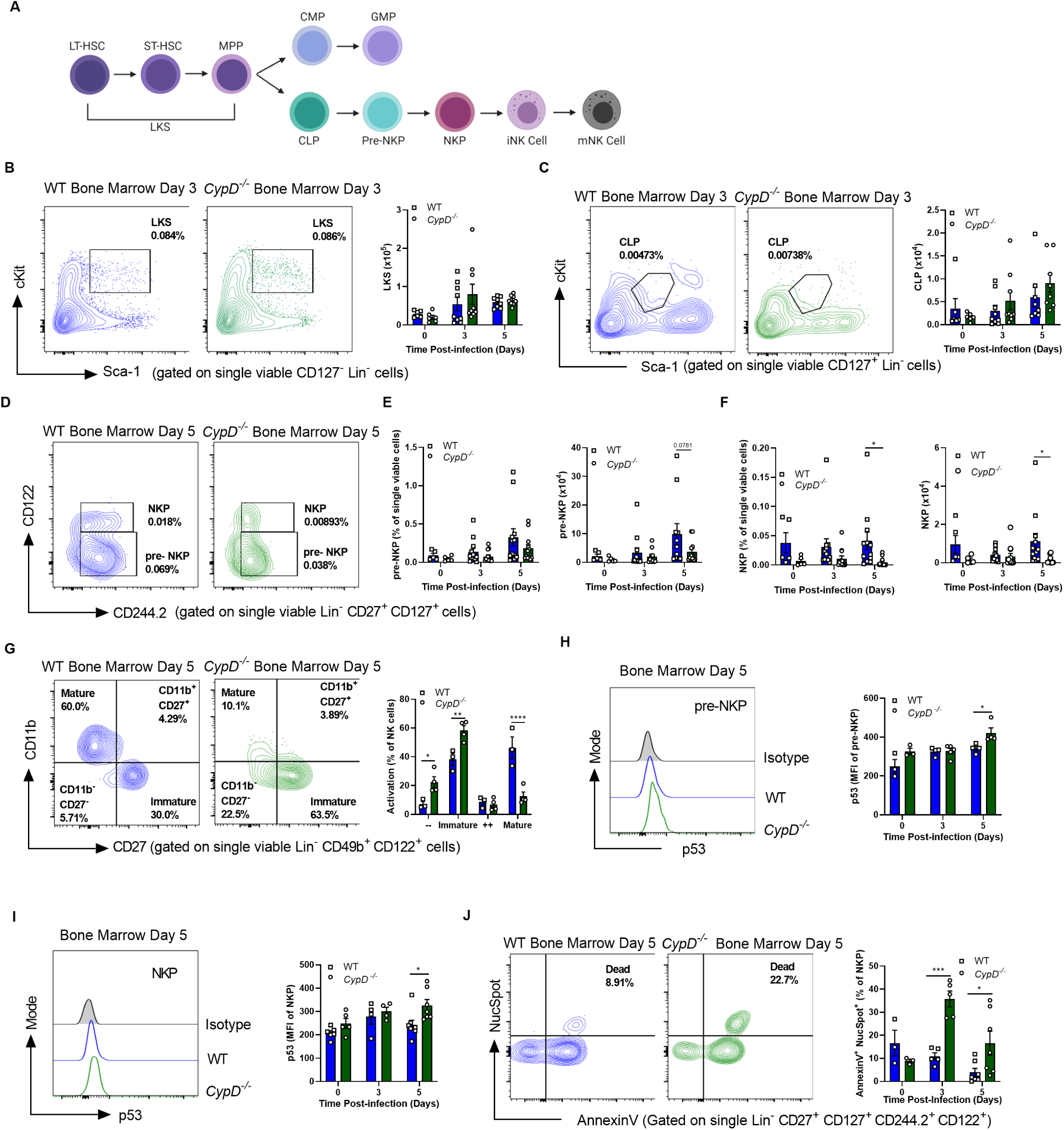
*CypD^-/-^* mice have reduced NK cell hematopoiesis in the bone marrow of infected mice due to cell death of progenitors. (A) Schematic of NK cell hematopoiesis starting at pluripotent LKS cells down to NK cells. (B-G) Total cell counts of LKS (B), CLP (C), pre-NKP (D-E), NKP (D, F) and differential activation statuses of mature NK cells (G) as assessed by flow cytometry. (D) Representative FACS plots of pre-NKP and NKP populations in the bone marrow at 5 days post-infection. In B, C and G, left panels are representative FACS plots taken at day 3 (B-C) and day 5 (G) post-infection. (H-I) Expression of p53 within pre-NKP (H) and NKP (I) populations in the bone marrow. Left panels are representative histograms taken at 5 days post-infection as quantified in the right panels. (J) Differential expression of AnnexinV and NucSpot to determine the levels of dead cells within the NKP population at various times post-infection. FACS plots on the left are taken from day 5 post-IAV infection. Panels in B-F and I-J are a compilation of at least 2 experiments, while G-H are one representative experiment of three independent experiments, with each symbol representing an individual mouse. In all panels, significance was assessed by Two-way ANOVA followed by Sidak’s multiple comparisons test. See also **Figure S4**.

### *CypD^-/-^* NK cell progenitors express higher levels of p53 and undergo enhanced cell death in the BM

To investigate the potential mechanisms involved in the reduction of NK cell progenitors in the BM of *CypD^-/-^* mice, we evaluated both cellular proliferation and death (*55, 56*). Both at steady-state and upon infection, CypD had no effect on proliferation of NK cell progenitors, as determined by intracellular Ki67 expression (**Fig. S4K-L**). Interestingly, p53, a central regulator of cell growth arrest and cell death in a variety of cell types (*57*), has recently been demonstrated to interact with CypD in mitochondria to facilitate necrosis of structural cells (*58*), while also antagonizing p53-dependent growth arrest in a tumour model, suggesting that CypD-deficiency could increase p53 function (*59*). Moreover, enhanced expression or activity of p53 halts lymphopoiesis, causes cell death specifically in the lymphoid lineage and leads to lymphopenia (*60, 61*). We, thus, investigated the level of p53 in NK cell progenitors by flow cytometry and found elevated p53 protein in *CypD^-/-^* pre-NKPs (**Fig. 6H**) and NKPs (**Fig. 6I**), but not in effector NK cells in the BM (**Fig. S4M**) or BAL (**Fig. S4N**). Using differential expression of AnnexinV/NucSpot (a 7-AAD analogue), the upregulation of p53 correlated with an enhanced percentage of death (AnnexinV^+^ NucSpot^+^) in NKP cells (**Fig. 6J**), but not in pre-NKP cells (**Fig. S4O**). Collectively, our results suggest that CypD antagonizes p53 function and prevents p53- mediated cell death to conserve NK cell lymphopoiesis and the generation of mature effector NK cells.

### Mice deficient in CypD are more susceptible to IAV due to a lack of IL-22 production by conventional NK cells

Having established that CypD regulates NK cell hematopoiesis and that *CypD^-/-^* NK cells are more phenotypically immature with altered transcriptomic and metabolic profiles, we finally hypothesized that NK cells from these mice are functionally impaired and confer susceptibility to IAV infection. As our RNA-seq data indicated impaired anti-inflammatory cytokine production (e.g. IL-10) and wound healing/hemostasis pathways in CypD-deficient NK cells (**Fig. 4B-C**), we speculated that altered cytokine production was responsible for the heightened susceptibility and pulmonary tissue damage.

Upon IAV infection, NK cells are a well-known source of IFN-γ (*62, 63*). However, the function of IFN-γ in response to IAV is controversial (*64–66*) and its potential role in disease tolerance has not been studied. At day 7 post-IAV infection, we observed a significant decrease in the levels of IFN-γ in the BAL of *CypD^-/-^* mice, which aligns with the increased pathology (**Fig. 7A**). Utilizing *ex vivo* stimulation of cells with PMA/ionomycin, we confirmed by intracellular cytokine staining (ICS) that CD49b^+^ NK cells from the BAL and spleen were substantial sources of IFN-γ. At day 5 post-IAV infection, a significantly lower frequency of *CypD^-/-^* NK cells was IFN-γ^+^ (**Fig. 7B; Fig. S5A**). However, in line with a previous study (*66*), using *Ifngr^-/-^* mice, we found no role for IFN-γ signaling in disease tolerance to IAV, with mice exhibiting similar amounts of protein and number of erythrocytes in the BAL (**Fig. S5B-C**), although a significant increase in total leukocytes was noted (**Fig. S5D**). To completely rule-out that the reduction in IFN-γ was responsible for the damage in the CypD-deficient mice, we reconstituted the airways of WT and *CypD^-/-^* mice with 100ng of IFN-γ intranasally (i.n.), or vehicle control, at 5 days p.i., collected the BAL at 7 days p.i and characterized pulmonary damage. As expected, we observed a significant increase in protein and erythrocytes in the BAL of CypD-deficient mice that received PBS compared to WT mice, but there was no amelioration in either group that received IFN-γ (**Fig S5E-F**). Collectively, these data confirm that, despite a reduction of NK cell-derived IFN-γ in CypD-deficient airways post-IAV infection, IFN-γ was not responsible for the break in disease tolerance.

**Figure 7:**
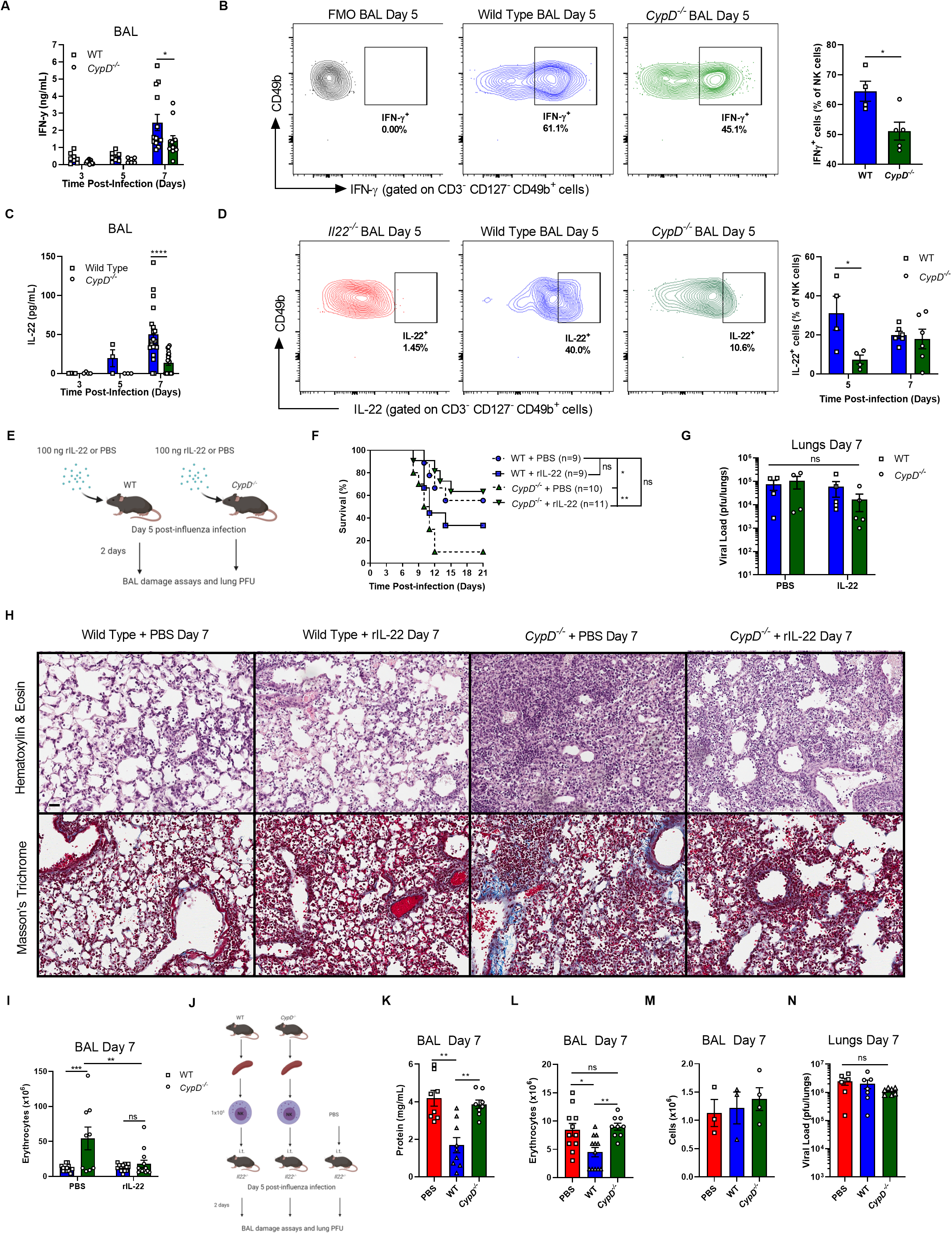
Reduced IL-22 production by NK cells in the airways is responsible for the susceptibility of *CypD^-/-^* mice. (A) Levels of IFN-γ in the BAL of infected WT and *CypD^-/-^* mice as determined by ELISA. (B) Intracellular cytokine staining of IFN-γ in NK cells. FACS plots are representative of the quantification on the right and are gated against the FMO. (C) Levels of IL-22 in the BAL of infected WT and *CypD^-/-^* mice as determined by ELISA. (D) Intracellular cytokine staining for IL-22 by NK cells in the BAL. FACS plots on the left are representative of the quantification on the right. An IL-22-deficient mouse was used as a staining control for specificity. (E) Model of recombinant IL-22 experiments as used in F-I. (F) Mice were infected with LD_50_ 90 PFU, administered recombinant IL-22 or PBS on day 5 p.i. and survival monitored. (G-I) a sublethal dosage of 50 PFU was used and mice were administered IL-22 or PBS as before. At day 7, viral loads were quantified (G), pulmonary inflammation and fibrosis were assessed by hematoxylin and eosin or Masson’s Trichrome staining, respectively (H; scale bar = 30µM) or erythrocytes in the BAL were enumerated (I). (J) Schematic of NK cell transfer experiments performed in K-N. On day 7 post-infection, day 2 post-transfer, protein (L), erythrocyte (M) and leukocyte levels in the BAL or lung viral loads (N) were assessed. In panels A, C, F, I, K, L and N data are combinations of two or three independent experiments. In B, D, G and M data are from one experiment that is representative of two or three independent experiments. Micrographs in H are representative of 3 or 4 individual mice. In A-D, G, I and K-N, each symbol represents data from one individual mouse, while in H total n values are displayed in the panel legend. Statistical analyses were performed as follows: in A-D Two-way ANOVA followed by Sidak’s multiple comparisons test; F Log Rank Test; G, I Two-way ANOVA followed by Tukey’s multiple comparisons test; K-N One-way ANOVA followed by Dunnett’s multiple comparisons test, Kruskal-Wallis test. See also **Figure S5**.

The production of IL-22 by NK cells plays a key role in protection of barrier sites through the promotion of epithelial cell survival and proliferation, and is a correlate of immunity to IAV (*29, 30*). Importantly, CD27^-^ NK cells, which are reduced in the BAL of *CypD^-/-^* mice, have been shown to be the major source of IL-22 with NK cell subpopulations (*28*). Thus, we postulated that a lack of IL-22 production by CypD-deficient NK cells was underlying their susceptibility to IAV infection. To assess this hypothesis, we began by confirming a previous study (*30*), showing that IL-22-deficient mice had enhanced lung damage, indicated by elevated protein and cell accumulation in the BAL, as well as pulmonary inflammation by histology at 7 days post-IAV infection (**Fig. S5G-I**). Next, we measured the level of IL-22 in the BAL of WT and *CypD*^-/-^ mice at various time points p.i. and found a significant reduction of IL-22 in *CypD^-/-^* mice at 7 days p.i. (**Fig. 7C**). Moreover, as with IFN-γ, we identified NK cells as an important source of IL-22 and there was a specific reduction of IL-22^+^ NK cells in both the BAL and spleen of CypD-deficient mice compared to WT (**Fig. 7D; Fig. S5J**). These data collectively confirm the importance of CypD within CD49b^+^ conventional NK cells in IL-22 production.

Because of the observation that IL-22 is involved in disease tolerance to IAV and that *CypD^-/-^* mice have reduced IL-22 in the airways, we investigated whether or not reconstitution of CypD-deficient mice with IL-22 could protect against IAV infection by limiting lung tissue damage. At 5 days post-lethal IAV infection, we intranasally reconstituted the airways of WT and *CypD^-/-^* mice with 100ng of IL-22 or PBS (**Fig. 7E**) and assessed mortality. As expected, *CypD^-/-^* mice that received PBS were significantly more susceptible than WT mice that received PBS. Strikingly, after IAV infection, the survival of CypD-deficient mice that received exogenous IL-22 was significantly increased and comparable to both WT groups (**Fig. 7F**). This enhanced survival was independent of host resistance (**Fig. 7G**), but dependent on disease tolerance as the increased levels of erythrocyte influx and fibrosis were abrogated in the CypD-deficient mice (**Fig. 7H-I**; **Fig. S5K**). Finally, having established the role of IL-22 in enhancing disease tolerance in *CypD^-/-^* mice, as well as an inability of CypD-deficient NK cells to produce IL-22, we sought to directly assess the capacity of WT versus *CypD^-/-^* NK cells to protect against tissue damage in an IL-22-deficient environment. To do this, we purified NK cells from naïve spleens of WT and CypD-deficient hosts, then transferred them (1×10^5^ cells) intratracheally into infected *Il22^-/-^* hosts at 5 days post-IAV infection, as to not affect early viral replication or the onset of tissue damage. At 7 days p.i. (2 days post-transfer), we collected the BAL and lungs, then assayed damage and viral loads (**Fig. 7J**). We found that IL-22-deficient mice that received WT NK cells exhibited statistically significantly enhanced disease tolerance, as assessed by attenuated protein and erythrocyte accumulation in the BAL, while no improvement was noted in mice receiving *CypD^-/-^* NK cells compared to PBS controls (**Fig. 7K-L**). In line with IL-22 acting primarily on structural cells, no variance in total leukocyte counts was observed in any group (**Fig. 7M**). Furthermore, no differences in viral load between any group could be delineated; thus, disease tolerance, rather than host resistance, was responsible for the reduced airway damage (**Fig. 7N**). Collectively, these data indicate that NK cell-derived IL-22 is dependent upon *CypD* expression and is required for regulating disease tolerance following IAV infection.

## DISCUSSION

The immune response to influenza requires a tightly regulated effort from both the innate and adaptive branches. Shortly following infection, early innate cells infiltrate the airways, and along with resident immune cells, coordinate to inhibit IAV replication and elimination (host resistance) from the lung. These host resistance mechanisms to IAV are well studied (*67*) and current influenza therapies, such as oseltamivir, exclusively target viral replication/dissemination to assist these pathways (*68*). However, host resistance comes at a substantial immunopathological cost that often is the cause of disease severity and mortality; therefore, limiting tissue damage and maintaining the functional capacity of the lung (disease tolerance) is critical for host survival. Breaks in disease tolerance cause mortality through loss of the integrity of the pulmonary epithelial/endothelial barrier, hemorrhaging and airway remodelling, all of which ultimately compromise gas exchange and lung physiology. Thus, the regulatory mechanisms of host defense against infection require a tight balance of generating immunity to resist a pathogen, while not compromising the physiology of infected organs that must be returned to homeostasis.

Disease tolerance as a defense strategy was initially established in plants (*69*) and has since been extended to mammals (*1, 3*). Evidence for its importance during chronic disease such as *Mycobacterium tuberculosis* and helminthic infections (*70, 71*) is accumulating, along with an understanding of the underlying mechanisms. Yet, our knowledge of disease tolerance pathways against acute viral infections like IAV remains limited. We and others have recently provided some insight into the importance of maintaining tissue integrity in disease tolerance to IAV either through the production of the host lipid mediator LTB_4_ (*16*) or inhibition of pulmonary matrix metalloproteases (*17*). In the current study, we extend the fundamental observation that reducing tissue damage is essential in disease tolerance and we define an role for CypD in this process, namely, by regulating NK cell generation in the bone marrow as well as promoting their effector functions in the airways. This work adds further evidence to our earlier studies that highlighted CypD as an essential component of lymphocyte-mediated host defense during infection (*72, 73*), in addition to its well-documented roles in necrosis (*74*) and the progression of neurological disorders, including multiple sclerosis (*75*).

The unique location of CypD within the mitochondrial matrix places it at the intersection of several essential cellular processes during health and disease. For example, in mature leukocytes, the mitochondrial antiviral signaling protein (MAVS) is anchored to the outer mitochondrial membrane and is essential for the induction of IFN-I in response to several viruses, including IAV. Although activation of the cytosolic pattern recognition receptor (PRR) RIG-I by viral RNA is the major trigger of MAVS-mediated responses, mitochondrial heat shock proteins, fission proteins and mitochondria-derived ROS critically regulate MAVS function (*76*). Additionally, IAV-encoded PB1-F2 localizes to the mitochondria and induces early intrinsic apoptosis to temper antiviral responses that is neutralized by the host NOD-like receptor NLRX1 (Nucleotide-binding, leucine-rich repeat containing X1) within the mitochondrial matrix (*35*). During LCMV infection, enhanced mitochondrial oxidative phosphorylation promotes antiviral immunity by pDCs (*77*), while our recent study highlighted the importance of CypD-dependent metabolic changes in T cell responses to promote disease tolerance to *Mtb* (*72*). Thus, mitochondria are critical orchestrators of both resistance and tolerance mechanisms in ways that are cell- and pathogen-specific.

Although the majority of previous work has focused on mitochondria in mature leukocytes, this study also revealed a role for mitochondrial CypD in regulating lymphopoiesis and p53- associated progenitor cell death. In structural cells, p53, an essential regulator of cell survival and death, was previously shown to translocate to the mitochondrial matrix and interact with CypD, resulting in necrotic cell death (*58*). Interestingly, enhancement of p53 activity specifically within lymphocyte progenitors has been shown to induce cell death and skew hematopoiesis towards myelopoiesis (*60, 61*). Our results uncovered a hitherto unappreciated inhibition of p53 by CypD in NK cell progenitors to promote lymphopoiesis, akin to the role previously described for Mysm1 and Mdm2 (*60, 61*). Importantly, we and others have described hematopoietic progenitor cell death as an emerging master regulator of bias between lymphopoiesis and myelopoiesis (*55, 78, 79*). Though no direct protein-protein interaction experiments between p53 and CypD were attempted in our study, it is intriguing to speculate that a balance between lymphopoiesis and myelopoiesis signals in progenitors may alter the outcome of p53/CypD interactions in the mitochondria, either by promoting or antagonizing p53-dependent lymphoid progenitor cell death as the leukocyte requirements change in the periphery. Certainly, more studies are required to fully delineate the contributions of CypD and p53 to hematopoiesis, as well as other known regulators such as Mysm1 and Mdm2.

In addition to our results in the bone marrow, we equally identified a function for CypD in mature NK cells that promotes their activation and function. NK cells are well known for their tumoricidal capacity and cytotoxicity against virally-infected cells, through an “innate” non-MHC restricted manner, which differs from cytolytic CD8 T cells (*19*). The importance of NK cells in the immune response to IAV is well-documented and appears to participate in both host resistance and disease tolerance. For example, complete genetic ablation of NK cells (via knockout of the transcription factor Nfil3; *Nfil3^-/-^* mice) (*22*), loss of the HA interacting receptor NKp46 (*Nkp46*^-/-^) (*23*), or depletion using an anti-asialo GM1 antibody (*24*) all result in IAV-induced lethality and elevated viral titres in the lung. Thus, NK cells are required for host resistance to IAV through HA/NKp46 cell lysis. While it has been recently shown that IL-22 production by NK cells limits pulmonary immunopathology during IAV infection (*29*), the regulatory mechanisms were unknown. Here, we found that CypD regulates IL-22 production by NK cells in the lung (**Fig. 7C-D**; **Fig. S5K**), while expression of perforin and granzyme B by NK cells (**Fig. 2H**) and viral load (**Fig. 1C**) were unaffected. Therefore, during the course of IAV infection, the functional capacity of NK cells is dynamic and can be changed from promoting resistance to tolerance.

Early NK cell responses, beginning at approximately 2 days post-IAV infection, appear to be mediated by NKp46 and are primarily concerned with restricting viral replication by killing infected cells, which correlates with the onset of symptoms. Thus, studies using mice that lack NK cells or NKp46 expression prior to the onset of infection show enhanced mortality coupled with increased viral loads (*22–24*). Following viral containment, NK cells may shift to a disease tolerance role that is marked by production of IL-22 beginning at around day 5 post-infection. Our results suggest that during the early phase of infection, the function of NK cells in host resistance is CypD-independent, while at the later stage of infection the function of NK cells in disease tolerance is mediated by CypD. As our RNA-Seq and functional data (**Fig. 3G**; **Fig. 3I-P**) highlighted elevated reliance on OXPHOS in *CypD^-/-^* NK cells compared to WT, we propose that a CypD-dependent metabolic shift in NK cells may, in part, dictate the transition of their effector program from host resistance to disease tolerance. Additional investigation is required to identify the intrinsic and/or extrinsic cues leading to the alteration in NK cell function during infection.

Greater consideration of the kinetics of infection may also serve to explain discrepancies in the source of IL-22 during IAV infection. IL-22 can be produced by several lymphocyte populations in response to IAV, including NK cells, NKT cells, ILCs and conventional αβ T-cells. Studies investigating the early phase of infection (i.e. 2 and 4 days p.i.) elucidated NKT and ILCs as the major sources (*26, 27*), while our and other’s work at the late phase of infection (i.e. day 5 and day 7) (*29*) showed conventional NK cells as the major source of IL-22. Collectively, these results suggest that early IL-22 production is dominated by NKT cells and ILCs, while NK cells are contributing to host resistance. However, at later stages of infection there is a shift in the function of NK cells highlighted by a production of IL-22, which contributes to disease tolerance. Although speculative, this concept is supported by our adoptive transfer experiments of NK cells into IL-22-deficient mice, where we show that CypD-dependent production of IL-22 by NK cells is sufficient to promote disease tolerance against IAV infection. Whether or not CypD solely regulates IL-22 production in NK cells rather than other cell types, or if its role is IAV-specific remains unknown. Undoubtedly, further insight into the kinetics of the source of IL-22 during IAV infection and how leukocyte function changes over time are required to more completely understand this complex phenomenon.

Our results outline an important role for CypD in NK cell development in the bone marrow as well as NK cell function in the airways. Although our work indicates separate roles for CypD in bone marrow NK cell progenitors and in mature NK cells, the exact mechanisms by which CypD functions and whether these two phenotypes are connected or separate require further investigation. The usage of a conditional knockout, rather than a constitutive knockout used in this study, that specifically deletes CypD in either mature NK cells or in bone marrow progenitors, could help to address these questions. Alternatively, mounting evidence suggests that progenitors may be imprinted in the bone marrow and this signature may affect the function of terminally differentiated effector cells in a process termed “trained immunity” (*80*). Thus, NK cells in the periphery of *CypD^-/-^* mice may be imprinted with an immature phenotype because of the lack of CypD function in bone marrow NK cell progenitors. We have recently described systemic BCG vaccination reprograms hematopoietic stem cells to promote myelopoiesis and generate more protective macrophage responses to subsequent *Mtb* infection (*79, 81*). Interestingly, NK cells have equally been shown to be “trained” by BCG (*82*), and memory NK cells are generated following IAV infection (*22*). Thus, it is possible that CypD may play a role in protective NK cell imprinting in the bone marrow. The potential of trained immunity during IAV infection is an exciting field for further research.

Influenza infections are persistent human pathogens, being responsible yearly for approximately 1 billion infections and between 300 000-500 000 deaths worldwide (*68*). Past IAV pandemics, as well as the current COVID-19 pandemic, have highlighted the importance of understanding host-pathogen interactions during acute respiratory infections. As we continue to better appreciate the interplay between host resistance (early responses) and disease tolerance (late responses), targeted therapies require an appropriate timeline for administration. For instance, our recent studies investigating host immunotherapy via the eicosanoid/IFN-I axis showed early inhibition of mPGES-1 (the enzyme involved in prostaglandin E_2_ production) increased antiviral IFN-β production to enhance host resistance (*83*). On the other hand, late-stage administration of exogenous LTB_4_ potentiates immunomodulation of IFN-α signaling to promote disease tolerance, without affecting antiviral responses (*16*). Considering that higher levels of lung immunopathology, rather than viral replication, is the major cause of morbidity and mortality during severe SARS-CoV-2 and IAV infections (*10, 84*), therapies that target disease tolerance may provide a more universal benefit compared to specific antiviral therapies. Studies that outline novel pathways of disease tolerance are required and will contribute to the next generation of respiratory virus therapies.

## MATERIALS AND METHODS

### Mice

Six- to ten-week-old C57BL/6, CD45.1 and *Ifngr^-/-^* mice were purchased from Jackson Laboratories. *CypD^−/−^* mice were provided by M. Forte (Oregon Health and Science University, Portland, OR, USA). All animals were housed and inbred at the animal facility of the Research Institute of McGill University. Experiments were performed using age- and sex-matched mice.

### Viruses & Infection

All *in vivo* infections were performed using mouse adapted influenza A/Puerto Rico/8/34 (H1N1) virus (IAV), kindly provided by Dr. Jonathan A. McCullers (St. Jude Children Research Hospital). Mice were challenged intranasally (in 25μL PBS) with IAV at a sublethal dose of 50 PFU or a lethal dose (LD_50_) of 90 PFU. 90 PFU was used for the survival experiments in **Fig. 1A** and **Fig. 7F**. For all other experiments, 50 PFU was used. During survival experiments mice were monitored twice daily for signs of duress and weighed daily. Mice reaching 75% of original body weight were considered moribund and sacrificed. Viruses were propagated and isolated from Madin-Darby Canine Kidney (MDCK) cells and titrated using standard MDCK plaque assays. MDCK cells were obtained from the American Type Culture Collection and maintained in Dulbecco’s modified Eagle medium enriched with 10% (*v/v*) FBS, 2 mM L-glutamine and 100 U ml^−1^ penicillin/streptomycin.

### Protein in the BAL

BAL were collected by cannulating the trachea with a 22-gauge cannula, then washing the lungs with 3 × 1 ml of cold, sterile PBS. The total volume recovered after lavage was ∼0.7 ml. Samples were spun down (1,500 r.p.m.; 10 min) and the total protein content was assessed by Pierce BCA Protein assay (Thermo Fisher Scientific).

### Texas Red-Dextran Lung Permeability

WT and *CypD^-/-^* were infected for 7 days with 50 PFU. On day 7, mice were delivered 25µL of 50mg/mL (1.25mg) Texas Red-Dextran (10 000 MW) intranasally. 1 hour later mice were sacrificed and lungs were carefully excised without damaging. Lungs were imaged using the In Vivo Xtreme (Bruker) using fluorescence capture. Resulting images were then analyzed for total fluorescent intensity of lung images using ImageJ software (National Institutes of Health). During the 1 hour of incubation Texas Red-Dextran molecules diffused into the blood of infected mice, due to the loss of epithelial/endothelial barrier integrity. Therefore, lower fluorescence is indicative of increased damage and compromised barrier integrity.

### Generation of chimeric mice

CD45.1^+^ B6 mice or *CypD^-/-^* mice were lethally irradiated with 9 Gy following 3 days of antibiotic treatment (0.5g Enrofloxacin (Bayer) per litre of drinking water). 16 hours later, the BM compartment was reconstituted with 4×10^6^ nucleated cells from either CD45.1^+^ mice (*CypD^-/-^* recipient) or *CypD^-/-^* mice (CD45.1^+^ recipient) and antibiotic treatment was maintained for 2 additional weeks. Between 10-12 weeks post-injection, reconstitution was validated by flow cytometry and was >90%. Mice were then infected for downstream assays.

### Flow Cytometry

Lung tissues were perfused with 10 mL of PBS, harvested and minced before collagenase digestion (150 U mL^−1^) for 1 h at 37 °C. Lungs were passed on a 40 µm nylon mesh, and red blood cells were lysed. For bone marrow staining, cells were isolated following aseptic flushing of the tibiae and femurs, and red blood cells were lysed. BAL were collected as previously described, spun down and red blood cells lysed. Spleens were aseptically removed, crushed on a 40 µm nylon mesh, and red blood cells were lysed. Then total cell counts were determined with a hemocytometer. In some experiments BAL were counted prior to red blood cell lysis to enumerate erythrocyte influx into the airways and then red blood cells were lysed. For peripheral blood staining, the blood was collected by cardiac puncture in a BD Microtainer tube and stained extracellularly; red blood cells were then lysed.

Cells were initially stained with eFluor-506 viability dye in PBS (eBioscience; 20 min; 4 °C), washed and surface stained with anti-CD16/32 (BD Biosciences) in 0.5% BSA/PBS solution to block non-specific antibody interactions with Fc receptors (10 min; 4 °C). Cells were then surface stained with combinations of PE-CF594-conjugated anti-SiglecF, BUV395-conjugated anti-CD11b, PerCP-eFluor780- or AlexaFluor700-conjugated anti-Ly6G, fluorescein isothiocyanate (FITC)- or allophycocyanin (APC)-conjugated anti-Ly6C, APC-eFluor780-conjugated anti-F4/80, BV421-conjugated anti-CD11c, FITC- or BUV395-conjugated anti-CD45.2 or APC-conjugated anti-CD45.1, Pe-Cy7-conjugated anti-CD3, BV786-conjugated anti-CD127, APC- or BUV737- conjugated anti-NKp46, BV421-conjugated anti-CD49b, PE-conjugated anti-CD49a (all from BD BioScience, except anti-Ly6G from eBioscience), or PE-Cy7-conjugated (BD Biosciences) or FITC-conjugated (eBioscience) anti-CD27. Cells were then fixed with 1% PFA for 1 hour, washed and acquired in 0.5% BSA/PBS solution.

In some experiments, following extracellular staining, samples were stained intracellularly for Ki67, active Caspase 3, perforin, granzyme B or p53. For APC-conjugated anti-Ki67, or anti-perforin and PE-conjugated anti-active Caspase 3, or anti-granzyme B (all from BD Biosciences) cells were initially fixed and permeabilized using BD Cytofix/Cytoperm (BD Biosciences) for 30 minutes at 4° C and then stained for 1 hour. Cells were washed and acquired. For p53 staining, cells were initially fixed and permeabilized for 1 hour using the Foxp3/Transcription Factor Staining Buffer Set (eBioscience) and then stained using the PE-conjugated anti-p53 Set (BD Biosciences), according to the manufacturer’s instructions. The provided PE-conjugated isotype was used as a control.

In experiments involving bone marrow progenitors, cells were processed, counted, stained for viability and blocked as before. Cells were then stained with biotin conjugated anti-Ly6C/G, anti-CD5, anti-B220, anti-Ter119, anti-CD4 and anti-CD8 for 20 minutes at 4° C. Cells were then washed and stained with APC-Cy7-conjugated streptavidin, APC-conjugated anti-cKit, PE-Cy7-conjugated Sca-1 for LKS cells. For CMP/GMP experiments, cells were not blocked and instead FITC-conjugated anti-CD34 and PerCP-eFluor780-conjugated anti-CD16/32 (all from BD Biosciences, except anti-CD16/32 from eBioscience) were added to the previous cocktail. For NK cell progenitors, PE-CF594-conjugated anti-CD122 and BUV395-conjugated anti-CD244.2 were added along with anti-CD27 and anti-CD127 (BD Biosciences) as previously described.

Finally, for experiments involving intracellular cytokine staining (ICS), 2×10^6^ single splenocytes or BAL cells were incubated for 4 hours at 37° C in the presence of PMA/Ionomycin and Brefeldin A (Cell Activation Cocktail; BioLegend) or GolgiPlug control (BD Biosciences). Cells were then stained extracellularly, fixed and permeabilized using BD Cytofix/Cytoperm (BD Biosciences), before being stained intracellularly for PE-conjugated anti-IL-22 and APC-conjugated anti-IFN-γ (BD Biosciences).

Flow cytometry acquisition was performed using BD LSRFortessa X-20 (BD Biosciences) with FACSDiva Software version 8.0.1 (BD Biosciences). Analysis was performed using FlowJo software version 10 (Tree Star).

### Evaluation of mitochondrial fitness using Mitotracker Green/Orange and Mitosox Red

Single cell suspensions were stained with extracellular antibodies as described above and then with Mitotracker Green and Orange 150nM, or MitoSox Red 1μM (Invitrogen technologies) in PBS for 30 min at room temperature and then washed with PBS. For experiments involving mitochondrial potential, dysregulated mitochondria were Mitotracker Green^hi^ and Mitotracker Orange^lo^, while respiring mitochondria were considered as Mitotracker Green^hi^ and Mitotracker Orange^hi^ as previously described (*85*)

### Extracellular Flux Analysis

Real-time oxygen consumption rates (OCR) of purified splenic NK cells were measured in XF media (non-buffered DMEM containing 2mM L-glutamine, 25mM glucose and 1mM sodium pyruvate) using a Seahorse XFe 96 Analyzer (Agilent Technologies). For the mitochondrial stress test, mitochondrial inhibitors oligomycin, fluorocarbonyl cyanide phenylhydrazone (FCCP), antimycin A and rotenone were used, as per the manufacturer’s recommendations. Briefly, NK cells were seeded at a density of 200 000 cells per well and 3 basal measurements were taken. Following this, 2 consecutive measurements were taken following each injection of oligomycin, FCCP, and antimycin A with rotenone. All measurements were normalized to cell number using a crystal violet dye extraction assay. Oxygen consumption curves were generated using Wave Desktop 2.3 (Agilent Technologies). Basal OCR was calculated by subtracting measurement 7 (non-mitochondrial respiration) from measurement 1. Maximal respiration was calculated by subtracting measurement 7 (non-mitochondrial respiration) from measurement 5 and spare respiratory capacity was the difference between maximal respiration and basal rate.

### Adoptive transfer model

NK cells were purified from uninfected spleens of WT and *CypD^-/-^* mice using the EasySep Mouse NK Cell Isolation Kit (Stem Cell Technologies) according to the supplier’s recommendations. Sorted cells were counted, washed (cold sterile PBS) and normalized to 1×10^5^ cells/50µL of sterile PBS. Purity was verified by flow cytometry and purity was always over 85% NK cells prior to transfer. NK cells were then transferred into *Il22^-/-^* mice on day 5 post-infection (50 PFU) via the intratracheal route. 2 days later, BAL were harvested for lung damage assays and, lungs were harvested for viral load analysis or histology.

### IL-22 or IFN-γ treatment

Recombinant murine IL-22 or IFN-γ was purchased from Peprotech. Mice were intranasally infected with 50 PFU of IAV. On day 5 post-infection, mice were given either PBS, IL-22 or IFN-γ (both 100ng/25µL) intranasally. Mice were sacrificed on day 7 post-infection, and the lungs were harvested and processed to determine the pulmonary viral load, or the BAL collected for damage assays. In some experiments, mice were infected with 90 PFU and IL-22 was delivered as stated and survival was monitored.

### Histopathological analysis

Lungs were inflated with and fixed for 48 h in 10% formalin, then embedded in paraffin. Next, 5 μm sections were cut and stained with Haematoxylin and Eosin or Masson’s Trichrome. Slides were scanned at a resolution of 20× magnification and pictures were taken using a Leica Aperio slide scanner (Leica). Quantification of collagen-afflicted areas on Masson’s Trichrome stained slides was performed using ImageJ software (National Institutes of Health) as previously described (*86*).

### Total bioactive IFN-I assay

Secretion of total active IFN-I (both IFN-α and IFN-β) in cell culture supernatants was assessed using the B16-Blue IFN-α/-β reporter cell line for murine samples (from InvivoGen), according to the specifications of the manufacturer. B16 cells were maintained in RPMI supplemented with 10% (*v/v*) FBS, 2 mM L-glutamine and 100 U ml^−1^ penicillin/streptomycin.

### Cell death analysis

Lactate dehydrogenase release in the BAL of IAV-infected mice was quantified using the CytoTox 96 Non-Radioactive Cytotoxicity Assay (Promega), per the manufacturer’s recommendations. Dead cell levels *in vivo* were assessed using PE-AnnexinV (BioLegend) and NucSpot Far-Red (Biotium), according to the manufacturer’s instructions and unfixed cells were acquired immediately by flow cytometry.

### Wet-to-dry ratio

Lungs were harvested from naïve or IAV-infected mice (50 PFU; day 7 post-infection), and blood clots were carefully removed. Then, the lungs were weighed (wet weight) and dried in an oven (56 °C, 2 d; dry weight), and the dry weight was measured. Data are presented as the ratio of wet weight to dry weight.

### RNA isolation and reverse transcription quantitative PCR (qPCR)

RNA from purified NK cells was extracted using RNeasy Kit (Qiagen) according to the manufacturer’s instructions. Some 500 ng of RNA were reverse transcribed using the ABM 5X RT MasterMix (Applied Biological Materials), as directed by the manufacturer. Complementary DNA was generated by qPCR using BrightGreen SYBR Green (Applied Biological Materials). Cq values obtained on a CFX96 PCR System (Bio-Rad) were analysed using 2^-ΔCq^ formula normalizing target gene expression to *Gapdh*.

### Library preparation and RNA-sequencing

Bulk RNA was collected from purified, splenic NK cells from 5 WT (1 control, 4 IAV-infected) and 5 *CypD^-/-^* mice (1 control, 4 IAV-infected). Sequencing libraries were constructed using the the Illumina TruSeq protocol. Libraries were sequenced on an Illumina NovaSeq (paired-end 100 base pair) to an average depth of 42.6 million reads per sample. Adapter sequences and low-quality score bases were trimmed from reads using Trim Galore (v0.6.2, Cutadapt v2.2) (*87*) in paired-end mode (-q 20 --paired --phred33). Trimmed reads were pseudoaligned to the *Mus musculus* reference transcriptome (mm10.81, downloaded from Ensembl) using the quant function in kallisto (v0.43) (*88*) in paired-end mode (average of 24.7 million pseudoaligned reads per sample). Gene-level expression estimates accounting for the average transcript length across samples were calculated using the R (v3.6.3) package tximport (v1.14.2) (*89*).

### Modeling the effect of the *CypD^-/-^* genotype on gene expression

Expression data was filtered for protein-coding genes that were sufficiently expressed across all samples (median logCPM > 1), leaving 12,451 genes for downstream analysis. After removing non-coding and lowly-expressed genes, normalization factors to scale the raw library sizes were calculated using calcNormFactors in edgeR (v3.26.8) (*90*). The voomWithQualityWeights function in limma (v3.40.6) (*91*) was used to apply these size factors, estimate the mean-variance relationship, convert counts to logCPM values, and estimate sample-specific observational weights that take into account variation in sample quality.

The following nested linear model was used to identify genes for which expression levels are differentially-expressed between WT and *CypD^-/-^* mice within each condition:

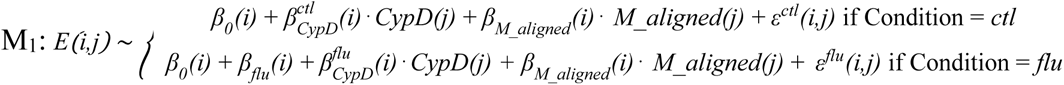

Here, *E(i,j)* represents the expression estimate of gene *i* for individual *j*, *β_0_(i)* is the global intercept accounting for the expected expression of gene *i* in a control WT mouse, 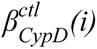 and 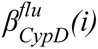 indicate the effects of the *CypD^-/-^* genotype on gene *i* within each condition, and *β_flu_ (i)* represents the intrinsic infection effect of IAV infection. Further, *M_aligned* represents the mean-centered, scaled (mean = 0, standard deviation = 1) number of pseudoaligned reads per sample (in millions), with *β_M_aligned_* being the impact of read depth on expression. Finally, *ε^cdt^* represents the residuals for each respective condition (*ctl* or *flu*) for each gene *i*, individual *j* pair. The model was fit using the lmFit and eBayes functions in limma, and the estimates of the genotype effect 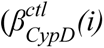 and 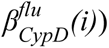 were extracted across all genes, along with their corresponding p-values. These estimates represent the genotype-related (WT vs *CypD^-/-^)* differential expression effects within each condition. We controlled for false discovery rate (FDR) using an approach analogous to that of Storey and Tibshirani (*92, 93*), which derives the null distribution empirically. To obtain a null, we performed 10 permutations, where genotype label (WT or *CypD^-/-^*) was permuted within infection condition (control or IAV-infected). We considered genes significantly differentially-expressed between genotypes if they had a 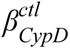 or 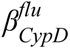 |logFC| > 0.5 and an FDR < 0.10. Of note, all of our downstream analyses focused only on the *CypD^-/-^* effect in the IAV-infected condition 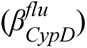.

### Gene set enrichment analyses

Gene set enrichment analysis was performed using two independent methods, fgsea (*94*) and ClueGO (*95*), depending on the type of data being evaluated. The enrichment program specifications and the data in which they were used to assess enrichments are described below:

The R package fgsea (v1.10.1) was used to perform gene set enrichment analysis for the genotype effects using the H hallmark gene sets (*96*). Input t-statistics were obtained directly from the topTable function in limma. These t-statistics were then ranked and used to perform the enrichment tests with the following parameters: minSize = 15, maxSize = 500, nperm = 100000. Enrichments scores (ES) and Benjamini-Hochberg adjusted p-values output by fgsea were collected for each tested gene set.

Gene set enrichment analysis was also performed for our lists of differentially-expressed (DE) genes between the WT and *CypD^-/-^* mice in the influenza-infected condition using the ClueGO (v2.5.7) (Bindea et al., 2009) Cytoscape (v3.7.1) (*97*) module in functional analysis mode, where the target set of genes was either the list of genes showing higher expression in the WT or *CypD^-/-^* mice and the background set was the list of all genes tested. Specifically, we tested for the enrichment of GO terms related to biological processes (ontology source: GO_BiologicalProcess-EBI-UniProt-GOA-ACAP-ARAP_08.05.2020_00h00) using the following parameters: visual style = Groups, default Network Specificity, no GO Term Fusion, min. GO Tree Interval level = 3, max. GO Tree Interval level = 8, min. number of genes = 2, min. percentage of genes = 4.0, statistical test used = Enrichment (right-sided hypergeometric test), p-value correction = Benjamini-Hochberg. For the graphical representation of the enrichment analysis, ClueGO clustering functionality was used (kappa threshold score for considering or rejecting term-to-term links set to 0.4). Only pathways with an FDR < 0.01 are reported.

To test for an enrichment of WT or *CypD^-/-^* DE genes in the IAV-infected condition among genes known to be involved in tissue repair and wound healing (TiRe gene set, curated from Yanai et al. (Yanai et al., 2016), n = 220 retained in our dataset), we calculated the proportion in our WT or *CypD^-/-^* DE gene lists that also fall into the TiRe gene set and considered this our “observed proportion”. To obtain a null distribution, we performed 1,000 permutations where, for each iteration, we: i) sampled the same number of genes as DE genes in our set (146 for WT and 169 for *CypD^-/-^*) from a list of all genes tested, and ii) calculated the proportion of these genes that are also in the TiRe gene set (our “null percentage”). P-values were computed by evaluating the number of permutations in which the null percentage was greater than or equal to the observed percentage divided by the number of total permutations (n = 1,000).

### ELISA

IFN-β levels in infected lungs and BAL were measured using a VeriKine Mouse IFN-β ELISA kit (PBL Assay Science). IFN-γ and IL-22 levels were assessed by ELISA (R&D Systems), as directed by the manufacturer.

### Statistical analysis

Data are presented as means ± s.e.m. Statistical analyses were performed using GraphPad Prism version 8.0.2 software (GraphPad). Statistical differences were determined using a two-sided log-rank test (survival studies), one-way analysis of variance (ANOVA) followed by Sidak’s multiple comparisons test, two-way ANOVA followed by Dunnett’s or Tukey’s multiple comparisons test, or two-tailed Student’s T-Test, as outlined in the Figure Legends. Significance is denoted by * p<0.05, ** p<0.01, ***p<0.001, ****p<0.0001.

### Ethics

All experiments involving animals were approved by McGill University (permit number 2010-5860) in strict accordance with the guidelines set out by the Canadian Council on Animal Care.

## AUTHOR CONTRIBUTIONS

M.D. conceived and supervised the project. J.D. and M.D. designed the experiments; J.D., E.P. and K.A.T performed experiments. Data were analyzed and interpreted by J.D. and M.D, except for RNA-seq data which were all analyzed and interpreted by H.E.R, under the supervision of L.B.B.. I.L.K. and S.A.K provided technical input/reagents, expertise and critically reviewed the manuscript. J.D., H.E.R., L.B.B and M.D. wrote the manuscript.

## DATA AND MATERIALS AVAILABILITY

All raw data generated for this paper are available at GEO GSE163290, and processed data files are accessible on Zenodo (doi: 10.5281/zenodo.4300712). All code used for analysis, along with associated documentation, is available at https://github.com/herandolph/CypD_flu.

## ACKNOWLEDGEMENTS

The authors would like to thank members of the Small Animal Imaging Labs (SAIL) of the RI-MUHC for assistance with the experiments in Fig. 1I, particularly Dr. Barry Bedell and Mathieu Simard. Additionally, the authors thank the Histopathology Core of the RI-MUHC for assistance with histology experiments, as well members of the Institut de recherches cliniques de Montréal (IRCM) for the generation and sequencing of RNA-seq libraries. Models in Fig. 6A, 7E and 7J were created by BioRender.com. This work was supported by the Canadian Institute of Health Research (CIHR) Project Grant (168885) to M.D.. M.D. holds a Fonds de recherche du Québec–Santé (FRQS) Award and the Strauss Chair in Respiratory Diseases. J.D. was supported by the Molson Foundation Award and RI-MUHC Studentship, H.E.R. was supported by a National Science Foundation Graduate Research Fellowship (DGE-1746045), E.P. was supported by a Fonds de Recherche du Québec–Santé Fellowship. The funders had no role in study design, data collection and analysis, decision to publish, or preparation of the manuscript.

## SUPPLEMENTARY FIGURE LEGENDS

**Supplementary Figure 1:**
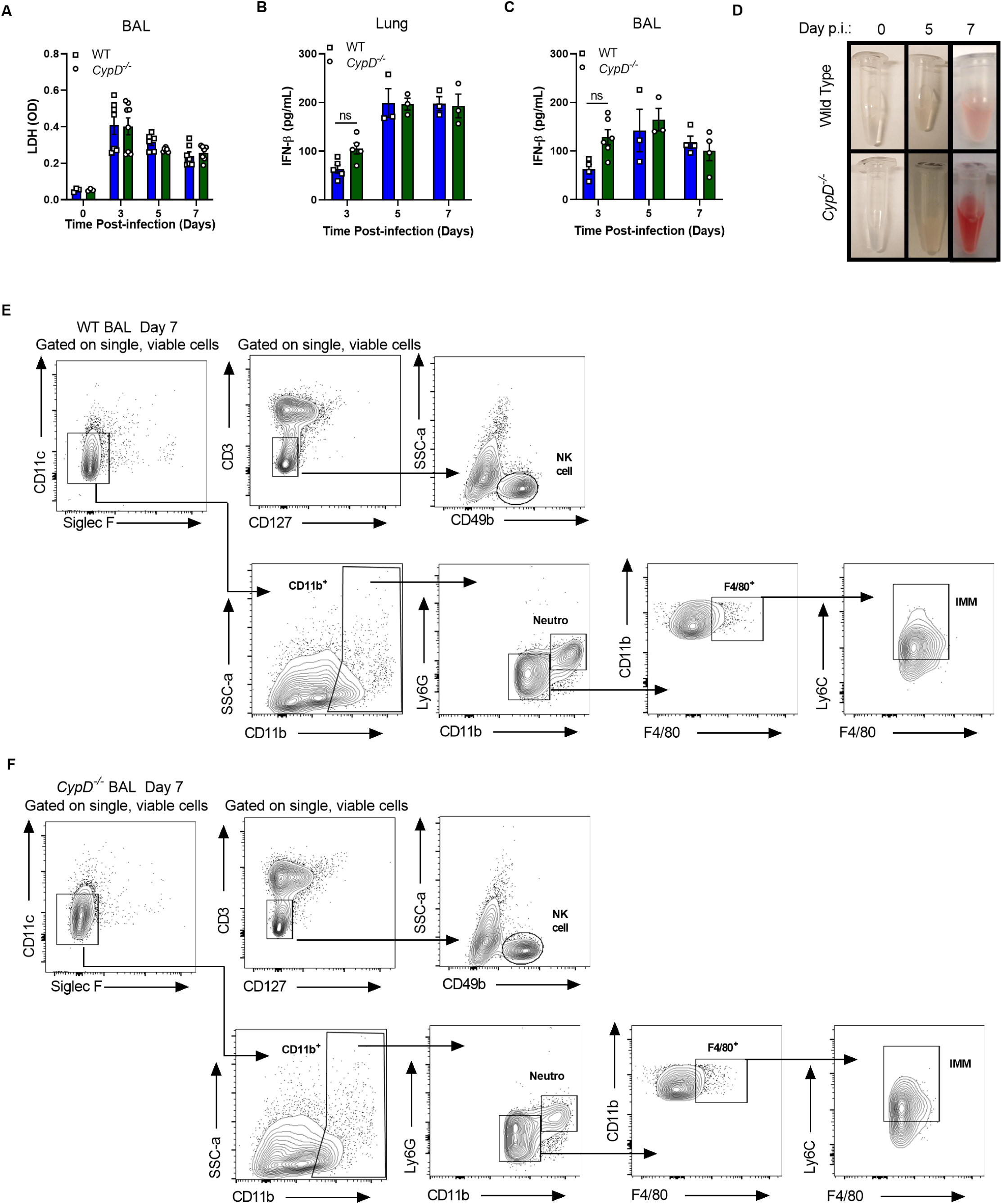
Susceptibility of CypD-deficient is due to disease tolerance and not host resistance mechanisms. (A-D) Mice were infected with 50 PFU of IAV and levels of LDH in the BAL (A), IFN-β in the lung (B) or BAL (C) were quantified. (D) Representative pictures of the BAL of mice over the course of infection as quantified in Figure 1H. Gating strategy for WT (E) and *CypD^-/-^* (F) mice used to quantify innate cells in the study. In panels A-C, each symbol indicates a separate mouse. Panel A is a compilation of two independent experiments and B-C are from one experiment. In panels A-C differences were assessed by Two-way ANOVA followed by Sidak’s multiple comparisons test. Refers to Figures 1-2

**Supplementary Figure 2:**
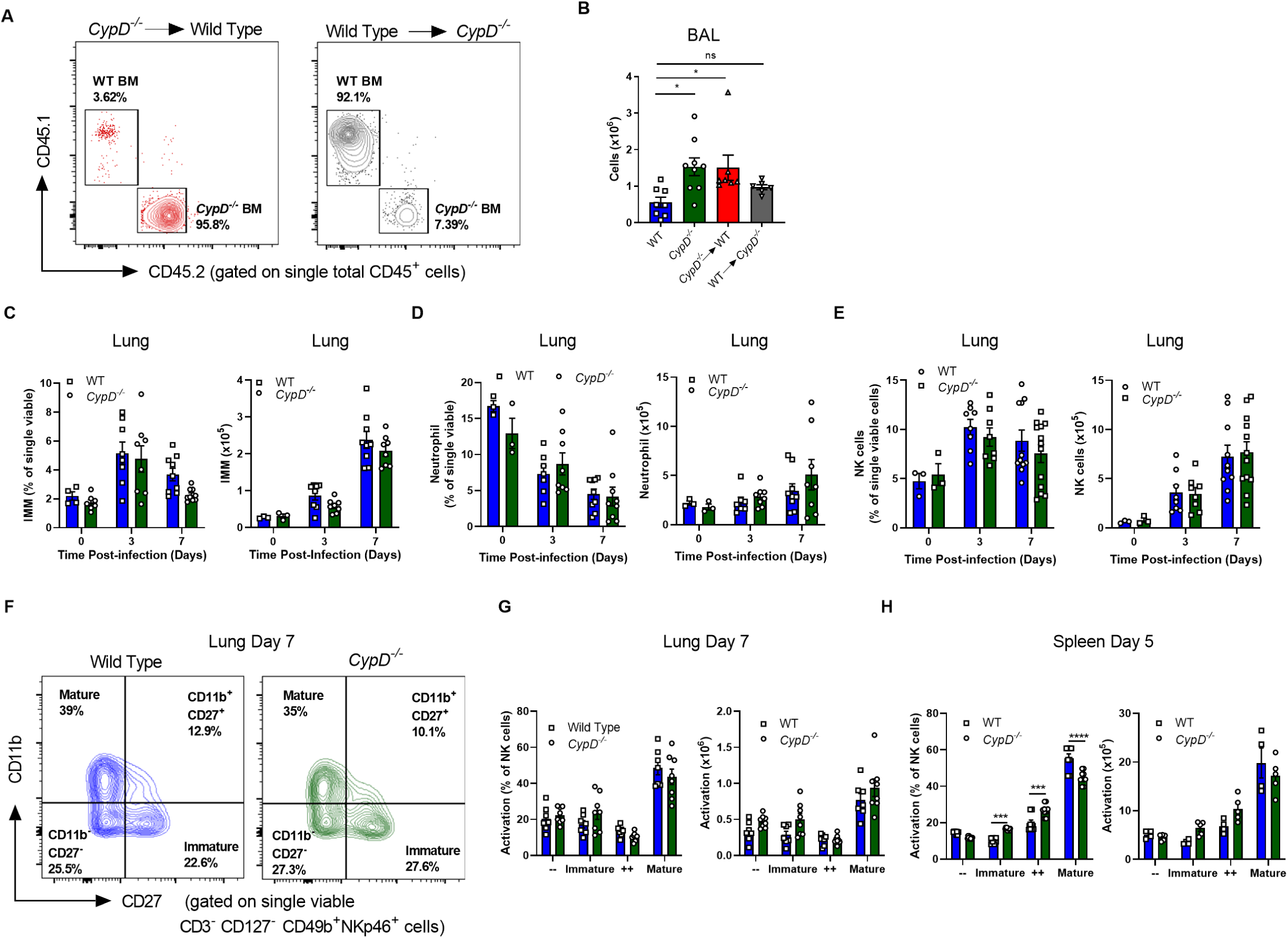
Kinetics and activation of NK cells. (A) Representative flow cytometry plots validating reconstitution of chimeric mice. (B-H) Mice were infected with 50 PFU of IAV. (B) Total cell counts in the BAL of chimeric mice at 7 days post-infection. (C-E) Frequencies (left panels) and total cell counts (right panels) of (IMM) (C), neutrophils (D) and NK cells (E) in the lung. (F) Representative FACS plots of NK cell activation in the lung at day 7 and quantified in (G) with frequencies in the left panel and total cell counts in the right. (H) Frequencies (left panels) and total cell counts (right panels) of NK cell activation in the spleen at day 5 post-IAV infection. In each panel, each symbol represents an individual mouse. Each panel is a compilation of two individual experiments, except H which is one representative experiment of two. Statistical differences were determined by One-way ANOVA followed by Dunnett’s test in B, or Two-way ANOVA followed by Sidak’s multiple comparisons test in all other panels. Refers to Figure 2.

**Supplementary Figure 3.**
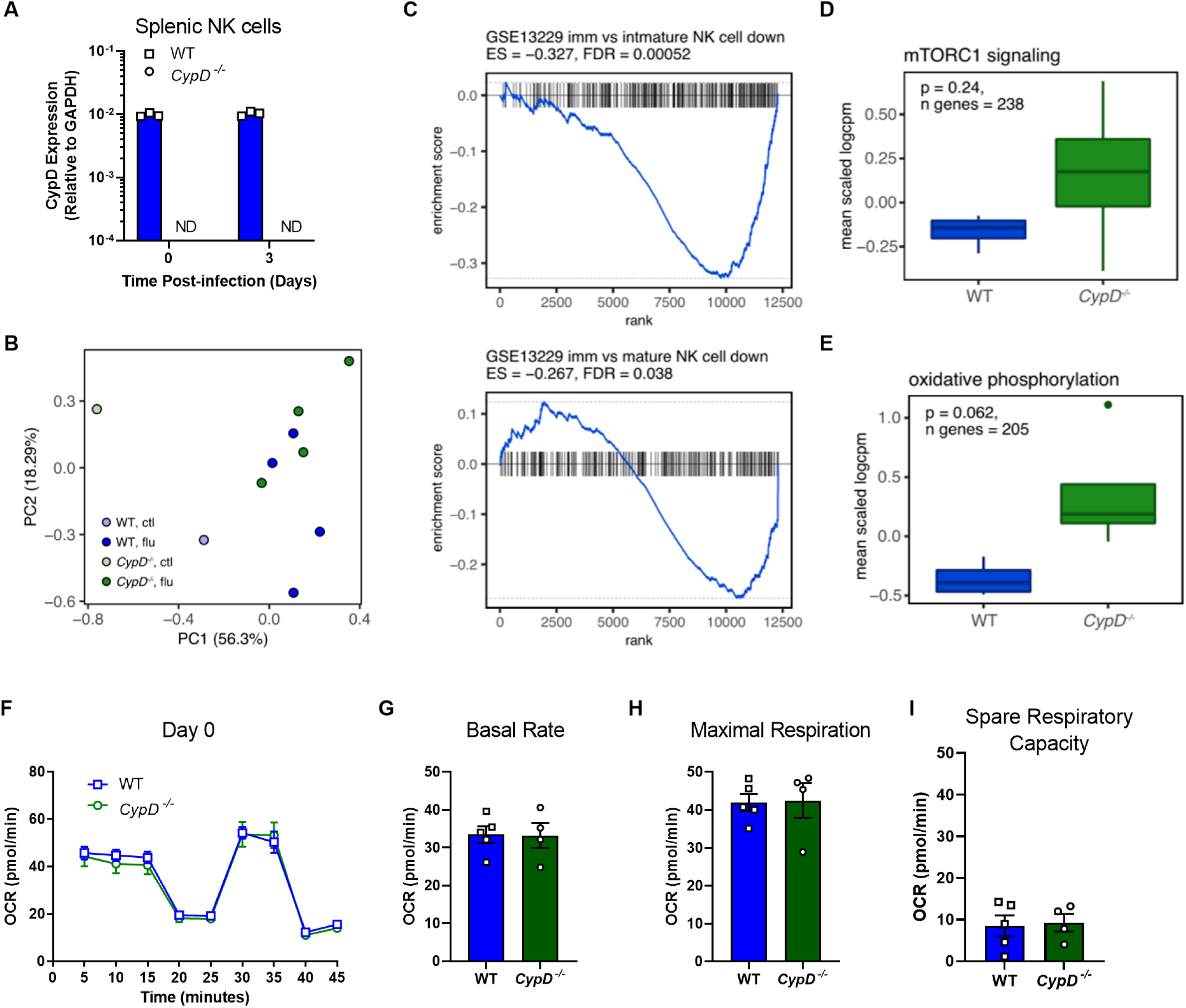
RNA-sequencing of purified, splenic NK cells in control and IAV-infected conditions. (A) Expression of *CypD* in purified NK cells by qPCR. (B) PCA decomposition of the splenic NK cell expression data in WT (blue) and *CypD^-/-^* (green) mice in control and IAV-infected conditions. PC1 (percent variance explained = 56.3%) separates samples by infection status. (C) Barcode enrichment plots for maturity marker gene sets as defined in Chiossone et al. 2009. A negative enrichment score (ES) corresponds to pathway enrichment among genes more highly expressed in WT mice. (D) Average, scaled logCPM expression estimates across genes in the mTORC1 signaling pathway and (E) the oxidative phosphorylation pathway in IAV-infected splenic NK cells of WT and *CypD*^-/-^ mice. (F-I) Oxygen consumption of naïve purified splenic NK cells as detailed in Figure 3I-P. Statistical differences were assessed in G-I by Student’s T-test. Refers to Figure 3.

**Supplementary Figure 4:**
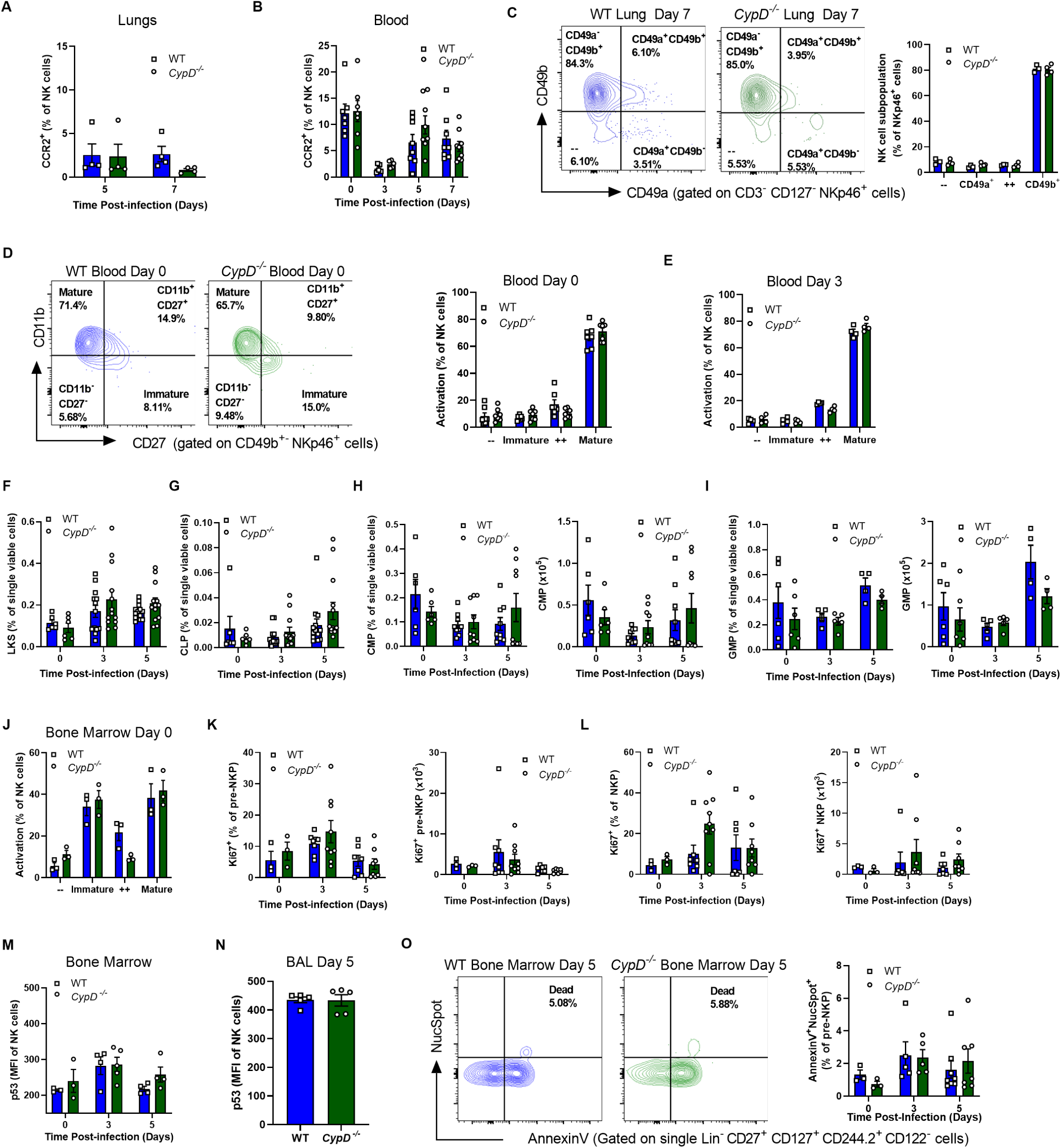
Kinetics of recruited NK cells and NK cell hematopoiesis in the bone marrow of WT and *CypD^-/-^* mice. (A-E) Mice were infected with 50 PFU of IAV. The percentage of CCR2-expressing NK cells in the lung (A) and blood (B). (C) Differential expression of CD49b and CD49a in the lung, with a representative FACS plot on the left and quantified on the right. (D-E) Activation status of NK cells in the blood in uninfected (D; representative FACS plot on the left) and day 3 (E) post-infection. (F-O) Mice were infected with 50 PFU of IAV and phenotyped by flow cytometry. Frequencies of LKS (F) and CLPs (G) following infection were determined as well as frequencies (left panels) and total cell counts (right panels) of CMPs (H) and GMPs (I). Relative activation states of naïve NK cells in the BM (J), as well as the percentage (left panels) and total cell counts (right panels) of Ki67-expressing pre-NKPs (K) and NKPs (L). Level of expression of p53 in NK cells in the bone marrow (M) and BAL (N) at day 5 post-infection. (O) Representative FACS plot (left panel) and quantification (right panel) of cell death in pre-NKPs as determined by AnnexinV and NucSpot staining. Symbols indicate an individual mouse and B, D, F-I, K-L and O are compilations of two individual experiments and A, C, E, J, M-N are one representative experiment of two. Statistical differences were assessed by Two-way ANOVA followed by Sidak’s multiple comparisons test for all panels, except for N where a Two-tailed Student’s T-test was performed. Refers to Figures 5 and 6.

**Supplementary Figure 5:**
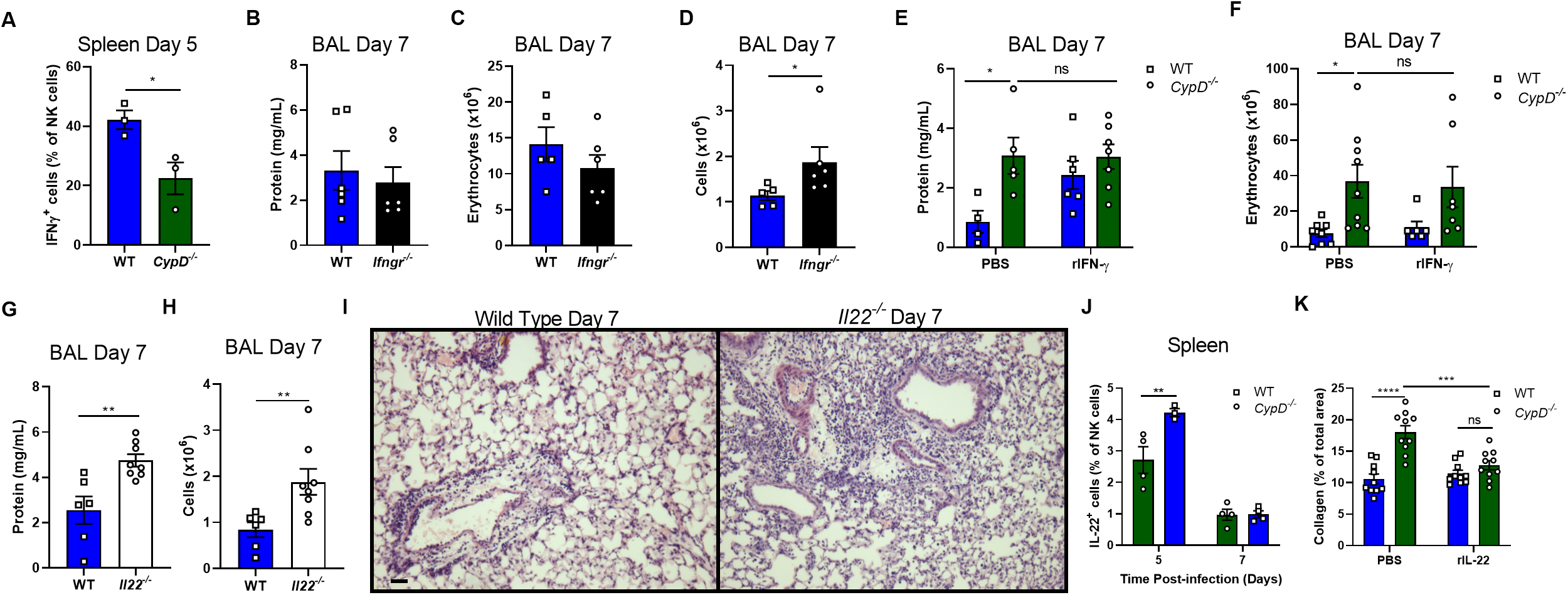
Disease tolerance is mediated by IL-22 and not IFN-γ following IAV infection. (A-K) Mice were infected with 50 PFU of IAV. (A) The frequency of IFN-γ-producing splenic NK cells at day 5 post-infection. (B-D) WT and *Ifngr^-/-^* mice were infected and the levels of protein (B), erythrocytes (C) and total cells (D) in the BAL at 7 days post-infection were determined. Following administration of recombinant IFN-γ, levels of protein (E) and erythrocytes (F) were assessed in the BAL. (G-I) WT and IL-22-deficient mice were infected and amount of protein (G) and number of cells (H) were enumerated, as well as pulmonary inflammation by hematoxylin and eosin staining (I; scale bar = 30µM). (J) Frequency of IL-22-producing NK cells in the spleens of WT and CypD-deficient mice at 5 and 7 days post-IAV infection. In all panels except I and K, symbols represent an individual mouse. In I, micrographs are a representative image taken from one of four mice. In K quantification is done from 10 random micrographs and each symbol represents one micrograph. In A and J, data are taken from one experiment that is representative of three. In B-H, panels are a compilation of two individual experiments. Differences were determined as follows: in A-D, G-H Two-tailed Student’s T-test; in E, F, K Two-way ANOVA followed by Tukey’s multiple comparisons test or Sidak’s (J). Refers to Figure 7.

Supplementary Table 1: RNA-Seq sample meta-data

Supplementary Table 2: CypD knockout effect on gene expression

Supplementary Table 3: Gene set enrichment analysis in WT and *CypD^-/-^* NK cells

